# Efficient genome-wide mapping of reproducible, context-dependent eQTLs at single-cell resolution

**DOI:** 10.64898/2026.08.25.747138

**Authors:** Jose Alquicira-Hernandez, Elizabeth Dorans, Yoshihiko Tomofuji, Aparna Nathan, Soumya Raychaudhuri

## Abstract

Single-cell technologies enable linking disease-risk variants to gene regulatory effects in specific cell-state contexts. However, most so called “single-cell eQTL” studies use a “pseudobulking” strategy to identify expression Quantitative Trait Loci (eQTLs), obscuring subtle dynamic regulatory effects of disease alleles. Here, we propose *Dynema* (**Dyn**amic **e**QTL **ma**pping in single cells) for fast and accurate genome-wide mapping of context-dependent and independent eQTL effects at true single-cell resolution. To identify eQTLs, *Dynema* uses a Poisson model with cluster robust variance estimators (CRVEs) to account for correlation of single-cell profiles from the same individual. In contrast to other common methods, *Dynema* achieves statistical calibration and scales to genome-wide analysis in large single-cell datasets in realistic timeframes. We applied *Dynema* to two independent T cell datasets and identified reproducible cell-state-dependent eQTL effects. Some cell-state-dependent eQTLs are missed by pseudobulking approaches, and many others are conditionally independent from lead eQTL effects. We show that *TSPAN32* and other autoimmune loci colocalize with cell-state-dependent eQTLs. Mapping context-dependent eQTLs at single-cell resolution enables the definition of the molecular effects of complex disease alleles.

## Main

Non-coding alleles drive ∼90% of complex disease loci^1^, and their molecular actions largely remain elusive. They likely alter gene regulation^2–4^, so they should induce gene expression changes as eQTLs; however, only ∼8% of GWAS loci colocalize with causal gene eQTLs^4,5^. This may be because disease alleles influence gene regulation specifically in critical cell states, while eQTLs are typically ascertained in heterogeneous samples, obscuring state-dependent regulatory effects. Characterizing the functional impact of disease variants requires methods for unbiased investigation of genetic effects on molecular phenotypes at granular cell-state resolution.

Single-cell eQTL analysis can link disease variants to cell-state-dependent gene expression variation. It can potentially characterize variable effects across healthy and disease states. We and others have proposed single-cell-resolution eQTL count-based models^6–8^, including a single-cell Poisson Mixed Effect (scPME) model, which we used to identify regulatory variant effects mediated by continuous T cell states and uncover autoimmune disease variants overlapping cell-state-dependent eQTLs.

There is still a critical need for statistically robust and scalable single-cell-resolution eQTL methods. First, accurately mapping eQTLs remains challenging due to sparsity and complex cellular heterogeneity. Gene expression distributions poorly fit analytical frameworks. Most methods rely on mixed-effect modeling to account for repeated donor measurements^6,8^ and cellular heterogeneity^6^. But mixed models make strong distributional assumptions and are sensitive to model misspecification^9,10^, resulting in inflated *p*-values and potential false positives^11^.

Second, the complexity of these models limits application to a small set of prioritized variants. Scaling single-cell eQTL mapping is commonly done by aggregation of single-cell profiles into “pseudobulk” profiles for analysis with bulk eQTL methods^12–16^. Single-cell models can be deployed to examine the “lead eQTL” variant to test for interaction effects with cell states^6,8,12^. But pseudobulk profiles may obscure disease-relevant biological signals. So, while practical, pseudobulk-first eQTL mapping misses potential state-specific regulatory variants that are independent of the lead eQTL in a locus.

To map single-cell resolution eQTLs genome-wide in large sample sizes at scale, we propose *Dynema*: a Poisson eQTL model with Cluster-Robust Variance Estimators (CRVEs) to account for cell correlation within donors.

## Results

### Overview of *Dynema*

*Dynema **(Dyn**amic **e**QTL **ma**pping for single cells*) maps eQTLs at single-cell resolution, scaling to genome-wide analyses in large datasets (**Methods**). Its flexible framework identifies main (i.e., context-independent) and interaction (i.e., dynamic context-dependent) eQTL effects, building on scPME^8^.

*Dynema* requires: a (1) single-cell expression matrix, (2) genotype matrix, (3) cell context matrix, and (4) matrix of cell- and sample-level covariates. Typical data includes thousands of cells per individual, thousands of genes, and high-density genome-wide genotyping. Contexts may be discrete (*e.g*. clusters) or continuous, reflecting dynamic cell processes. Continuous context may be functions of expression data or other data modalities, including latent spaces such as principal component analysis (PCA)^17^, canonical correlation analysis (CCA)^18^ or non-negative matrix factorization (NMF)^19^.

*Dynema* models expression counts with a Poisson Generalized Linear Model (GLM) as a function of variant allele dosage, context(s), and genotype x context interaction(s), controlling for sample and single-cell covariates (**Figure 1a**) (**Methods**). Individual genes have different underlying distributions (**Figure 1c**). *Dynema* uses cluster-robust variance estimators (CRVEs^20–22^, **Methods**). Unlike naïve variance estimation, which relies on theoretical distributional assumptions, CRVE measures residual variance empirically, thereby relaxing the Poisson mean-variance equality assumption and yielding valid estimates even when those assumptions are violated. CRVE accounts for within-donor correlation of single cells by grouping residual variance within each donor to produce robust standard errors (**Methods**). This is important because cells from the same donor may share biological or technical non-independent sources of variation.

**Figure 1.**
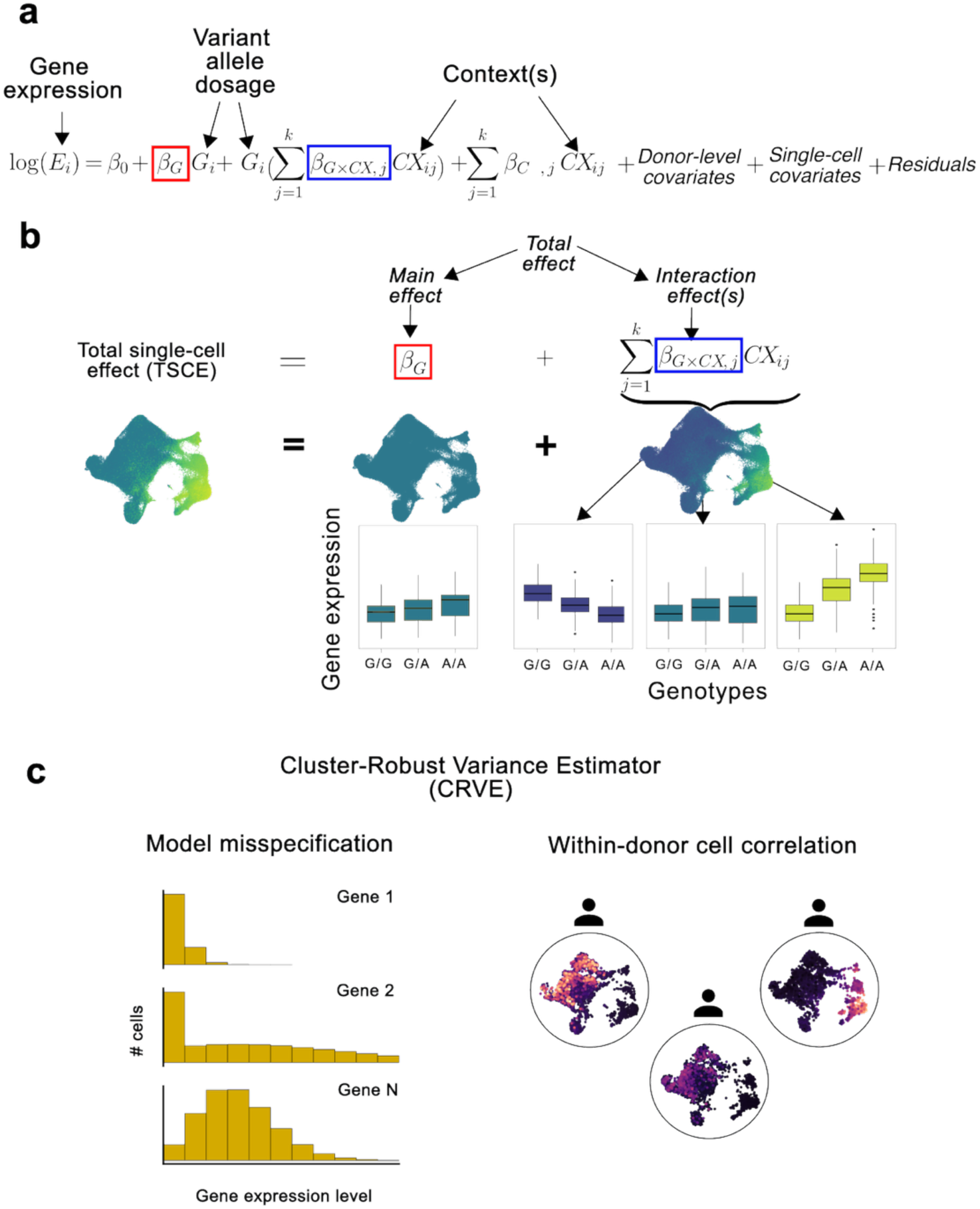
Schematic of *Dynema* framework. **a**, *Dynema* uses a Poisson GLM architecture for single-cell eQTL mapping of main, interaction, or total effects. *E* is a vector with length n containing the expression of the gene in each cell. *G* is the allele dosage of a given variant. Contexts (CX) are modeled as continuous or discrete numeric weights for each cell. Biological and technical covariates are also modeled at the donor and single-cell levels. **b**, Total, main, and interaction effects that *Dynema* can map. UMAPs show: (left) a total single-cell effect (TSCE), computed by adding the main effect *β_G_* and the single-cell level interaction effects, (middle) main effect, and (right) interaction effect(s). Boxplots show the effect of each allele on gene expression dependent on the context(s) **c,** Cluster-robust variance estimators (right) avoid model misspecification for different gene expression distributions and (left) capture within donor correlation of gene expression across cells.

While *Dynema* provides robust analytical *p*-values, it can also derive empirical *p*-values through score bootstrapping^20,21^ for extreme scenarios such as small sample size or skewed cell counts (**Methods**).

*Dynema* decomposes eQTL effects into: 1) a ***main effect***, independent of cellular context, capturing the average variant effect on expression, and 2) ***interaction effect(s)***, capturing context-specific effects (**Methods**). If multiple contexts are considered, we can assess the significance of each individual ***single-context interaction effect*** or of ***multi-context interaction effects*** in aggregate with a multiple degree-of-freedom test. *Dynema* can also test the ***total effect*** by combining main and interaction components (**Figure 1b**). We recommend first testing the total effect to identify eGenes, and then testing main and interaction effects separately to characterize context-independent and -dependent signals. For each cell, we can define a **total single-cell effect (TSCE)** by combining the main effect *β_G_* eQTL component and the interaction component, which we can calculate using cell state values and the genotype-cell-state interaction *β* values (**Figure 1b**).

### Genotyped single-cell datasets for testing

We used a memory T cell CITE-seq dataset^8^ (TBRU) comprising 259 donors and ∼500,000 cells. Previously, we annotated T cell states integrating single-cell RNA and protein surface abundance via CCA^13^ (**Supplementary Figure 1**). We used the top three canonical variates (CVs) capturing essential cellular contexts (i.e. cell states) and accounting for most of the variance in both modalities: cytotoxicity (CV1), regulatory/activation (CV2), and central memory (CV3) phenotypes^23–25^.

To assess cell-state-dependent eQTL reproducibility, we also analyzed ∼579,000 T cells from 969 donors in OneK1K^12^. Since OneK1K lacked surface protein expression, we could not directly calculate CVs. Instead, we inferred CV1-3 in OneK1K with Symphony^26^ (**Methods**). Inferred CVs capture similar biology as the original CVs, given the concordance of author-reported cell type labels, CV scores, and marker gene expression (**Supplementary Figure 2 and 3**).

To ensure power to detect dynamic eQTLs, we restricted our analysis to genes with non-zero expression in >50% of donors and >5% of cells, yielding 6,289 genes in TBRU and 3,821 in OneK1K, with 3,740 genes shared between them (**Supplementary Figure 4**).

### Dynema’s type I error is well-calibrated

We evaluated statistical inflation in *Dynema* and four other single-cell eQTL models for mapping cell-state-dependent eQTLs (CASTIE^27^, *scPME*^8^, *CellRegMap*^6^, single-cell linear mixed-effect model (*scLME*), **Methods**). Using TBRU, we obtained lead eQTL variants for 2,202 genes. We permuted genotypes to create null data, while preserving interindividual differences and cell-state expression structure. This scenario reflects the common case for variant-gene pairs, where no main or interaction effect exists.

When we tested single-context interactions with CV1 (GxCV1)*, Dynema* demonstrated well-calibrated type I error (**Figure 2a**). Analytical and bootstrap *p*-values were <0.01 in 0.8% and 0.7% of tests, respectively (**Figure 2b**), with observed *p*-values consistent with expectation under the null. CASTIE also demonstrated well-calibration with 1.09% significant tests at *p* < 0.01 but had several highly significant tests (*p* < 1 × 10^-5^, **Figure 2a**). In contrast, other methods showed clearly inflated *p*-values <0.01: scPME (2.6%), CellRegMap (4.6%), and scLME (3.9%) (**Figure 2a-b**), in some cases identifying highly significant false positive associations (*p* < 1 × 10^-5^, **Figure 2a**). We quantified inflation with an inflation factor *λ*, capturing systematic inflation of the median statistic (**Methods**). *Dynema* was well-calibrated (*λ_analytical_* = 1.04, *λ_boot_* = 1.04) as well as CASTIE (*λ* = 0.99), while other methods showed inflation (1.20 ≤ *λ* ≤ 1.41, **Figure 2c**). *Dynema* was also calibrated for multi-context interaction, main, and total effects (**Supplementary Note 1; Supplementary Figure 5-9**).

**Figure 2.**
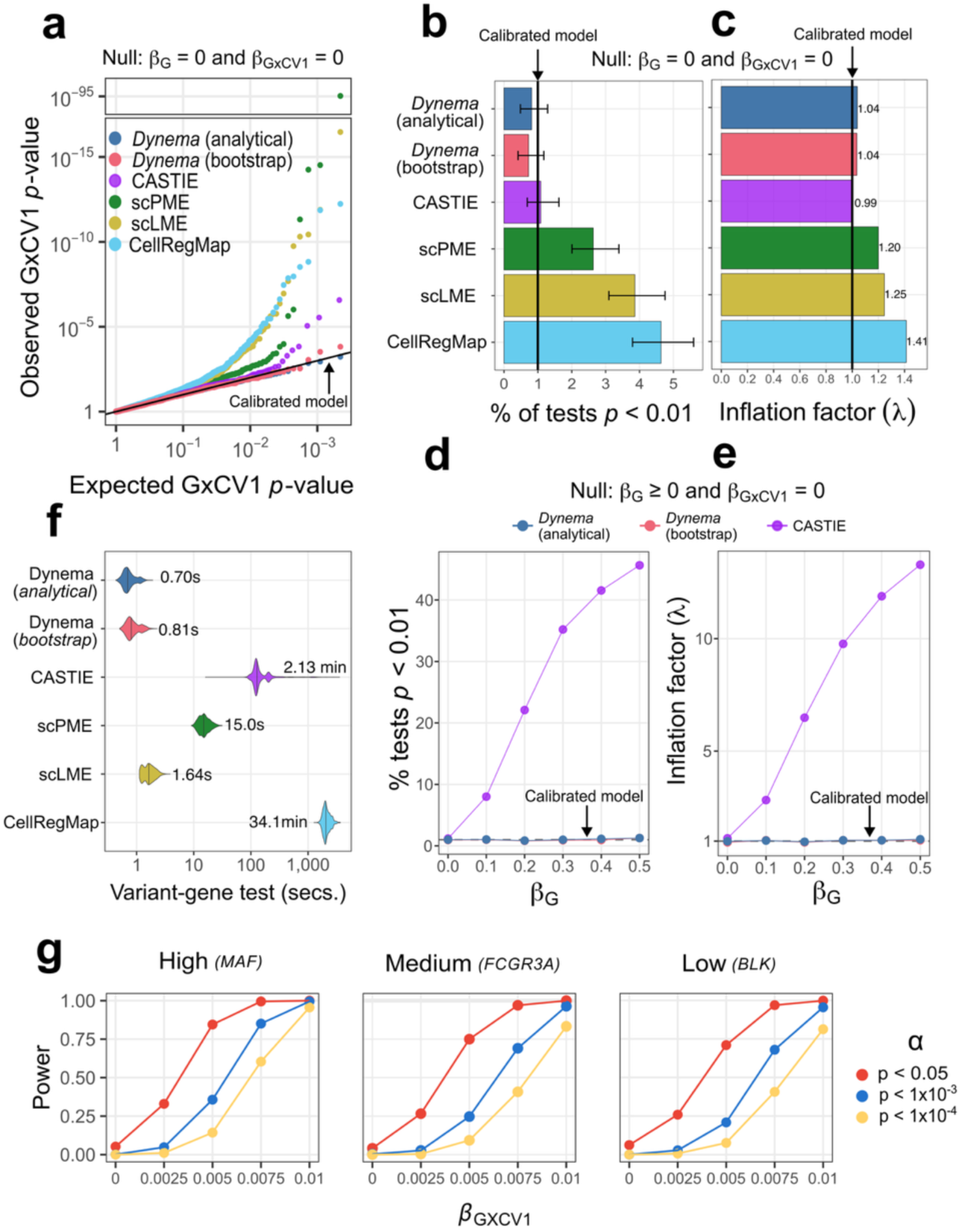
Benchmarking *Dynema*’s statistical calibration, speed, and power analysis. **a**, Q-Q plot showing the expected and observed p-values for the GxCV1 term in five single-cell eQTL mapping approaches. Expected *p*-values were computed under the null scenario generated by permuting genotypes: *β*_G_ = 0 and *β*_GxCV1_ = 0. Each point corresponds to one lead variant-gene test (2,202 genes in total). **b**, Bar plot of percent of significant tests at *p* < 0.01 for each test. Confidence intervals are from an exact binomial test. **c,** Inflation factor for *Dynema* and other methods calculated from **a**. **d**-**e,** The Y-axes represent the percent of significant tests at *p* < 0.01 and inflation factor λ, respectively, with null *β*_G_ ≥ 0 and *β*_GxCV1_ = 0 in parametric Poisson simulation. The X-axes shows the main-effect beta increasing from left to right. The dashed lines represent the expected % of significant events at the null (1%) in **d**, and λ=1 in **e**. **f**, Speed comparison of *Dynema* and other approaches. X-axis shows the elapsed time in seconds for a single variant-gene association test, and the number and horizontal line show the median across testing 2,202 lead variant-gene pairs for all methods, except CASTIE (n=2,122; **Methods**). **g**, *Dynema*’s statistical power for interaction eQTL effects with CV1 for high, medium and low expressed genes. Each panel shows a different gene representative of distinct levels of average mean expression (0.345, 0.054, 0.004 UMIs) in the TBRU dataset. The X-axis shows the interaction effect size (change in natural log mean expression per allele copy x CV1 unit) that was simulated using a Poisson GLM. Y-axis shows the proportion of true associations detected at distinct significance levels, each shown in a different color.

We tested more challenging and realistic scenarios where genuine main effects are present but context-dependent interaction is absent (*β_G_* > 0 and *β_CV1-3_* = 0). Using a simple parametric Poisson simulation (**Methods**), *Dynema* maintained robustly calibrated interaction *p*-values regardless of the main effect’s magnitude (**Figure 2d-e**). The percent of tests with *p* < 0.01 ranged from 0.86% to 1.27% (analytical) and 0.82% to 1.27% (bootstrap). The corresponding inflation factors (λ) ranged from 0.97 to 1.08 (analytical), and 0.94 to 1.04 (bootstrap). This was also true when testing multi-context interactions with CV1-3 and using a data-based simulation approach^28^ (**Methods; Supplementary Figure 10 and Supplementary Note 2**). Since CASTIE was well calibrated in the absence of main effects, we examined its performance too. CASTIE demonstrated severe statistical inflation in these scenarios, systematically misidentifying these common main effects as false-positive interaction signals with % of tests with *p*<0.01 from 8.01% (for very small main effects β_G_= 0.1) to >45% (with larger but realistic sized main effects) (**Figure 2d**) and λ ranging from 2.82 to 13.3 (**Figure 2e**).

This miscalibration behavior was also observed when testing multi-context interactions (CV1-3) and using a data-based simulation approach^28^ (**Methods; Supplementary Figure 10 and Supplementary Note 2**).

We tested *Dynema* in more extreme conditions with fewer individuals or highly variable cell counts (**Methods**). While analytical *p*-values showed some evidence of inflation in the most extreme situations (*λ* = 1.04-1.22; **Supplementary Figure 11a**), bootstrap *p*-values remained well-calibrated (*λ* = 0.99-1.05; **Supplementary Figure 11b**) in all scenarios. *Dynema* is generally robust to low sample sizes and extreme imbalances in cell counts; but bootstrapped *p*-values can be used for confirmation if necessary.

### *Dynema* scales efficiently to enable genome-wide analysis

Next, we assessed *Dynema*’s computation time on a standard CPU (**Figure 2d**). Dynema completed a multi-context eQTL interaction analysis for a single variant-gene pair in half a million cells in ∼0.7 seconds. Bootstrapping had a similarly fast computation time. It was on average 21.4 times faster than our previously proposed *scPME* model, and 182.9 and 2925.8 times faster than CASTIE and *CellRegMap*. With parallelization across 4 threads, *Dynema* completes a locus-wide analysis in < 1 minute.

### *Dynema* is well-powered to detect eQTL interactions with cell states

We assessed *Dynema*’s power to detect context-dependent regulatory effects. We selected three genes with high, medium, and low expression in TBRU (*MAF*, *FCGR3A*, *BLK*) and modeled data with average expression of 0.345, 0.054 and 0.004 UMIs per cell, respectively. We simulated eQTLs by varying GxCV1 interaction effect sizes. We evaluated *Dynema*’s power to detect the GxCV1 interaction at *α* = 0.05, 1 × 10^-3^, and 1 × 10^-4^ (**Figure 2e; Methods**). Realistic CV1 interaction effect sizes in TBRU ranged from 8.6 × 10^-5^ to 0.28 (mean = 0.044). *Dynema* showed strong power across these genes to detect realistic eQTL interactions. Additionally, *Dynema* identified main and interaction effects largely consistent with those reported for 6,511 genes in our previous study using scPME (see **Supplementary Note 3; Supplementary Figure 12)**.

### Many lead eQTL variants interact with cell states

We assessed lead main effect eQTL variants for cell-state-dependent eQTL effects. We pseudobulked cells and defined lead eQTLs independently in TBRU and OneK1K. To ensure high-confidence eQTLs, we assessed significance stringently correcting for the total variant-gene pairs tested (Šídák^29^, **Methods**). In TBRU and OneK1K, we identified 1,682 eQTLs across 6,289 genes (*p*<8.7 × 10^-9^; n=5,890,440 pairs; **Supplementary Table 1**) and 1,456 eQTLs across 3,821 genes (*p*<1.2 × 10^-8^; n=4,227,088 pairs; **Supplementary Table 2**) respectively.

*Dynema* can test *multi-context interaction effects* simultaneously. We tested lead eQTL variants for interaction effects with three CVs in aggregate, identifying dynamic eQTLs with false discovery rate (FDR)^28^ < 0.05. In TBRU and OneK1K, 631 (38%) and 513 (35%) lead variants interacted with cell states (**Supplementary Figure 13; Supplementary Table 3-4**), consistent with our previous study^8^.

### *Dynema* identifies reproducible eQTL interactions across datasets

We assessed *Dynema*’s ability to replicate interaction eQTL effects, focusing on lead main eQTLs in TBRU and OneK1K, that are not cohort or ancestry specific. We meta-analyzed pseudobulk eQTL profiles from TBRU and OneK1K to identify 1,719 shared variant-gene pairs (*p* < 1.7 × 10^-8^; n = 2,939,514) across 3,740 genes expressed in both datasets (**Supplementary Table 5; Methods**).

We applied *Dynema* to these 1,719 variant-gene pairs for main effects in TBRU and OneK1K without CV interactions (**Supplementary Tables 6-7**). Unsurprisingly, allelic direction (concordance = 0.99, **Figure 3a**) and z-scores were concordant (*r* = 0.89) for nominally significant main effects (*p* < 0.05 in both datasets),

**Figure 3.**
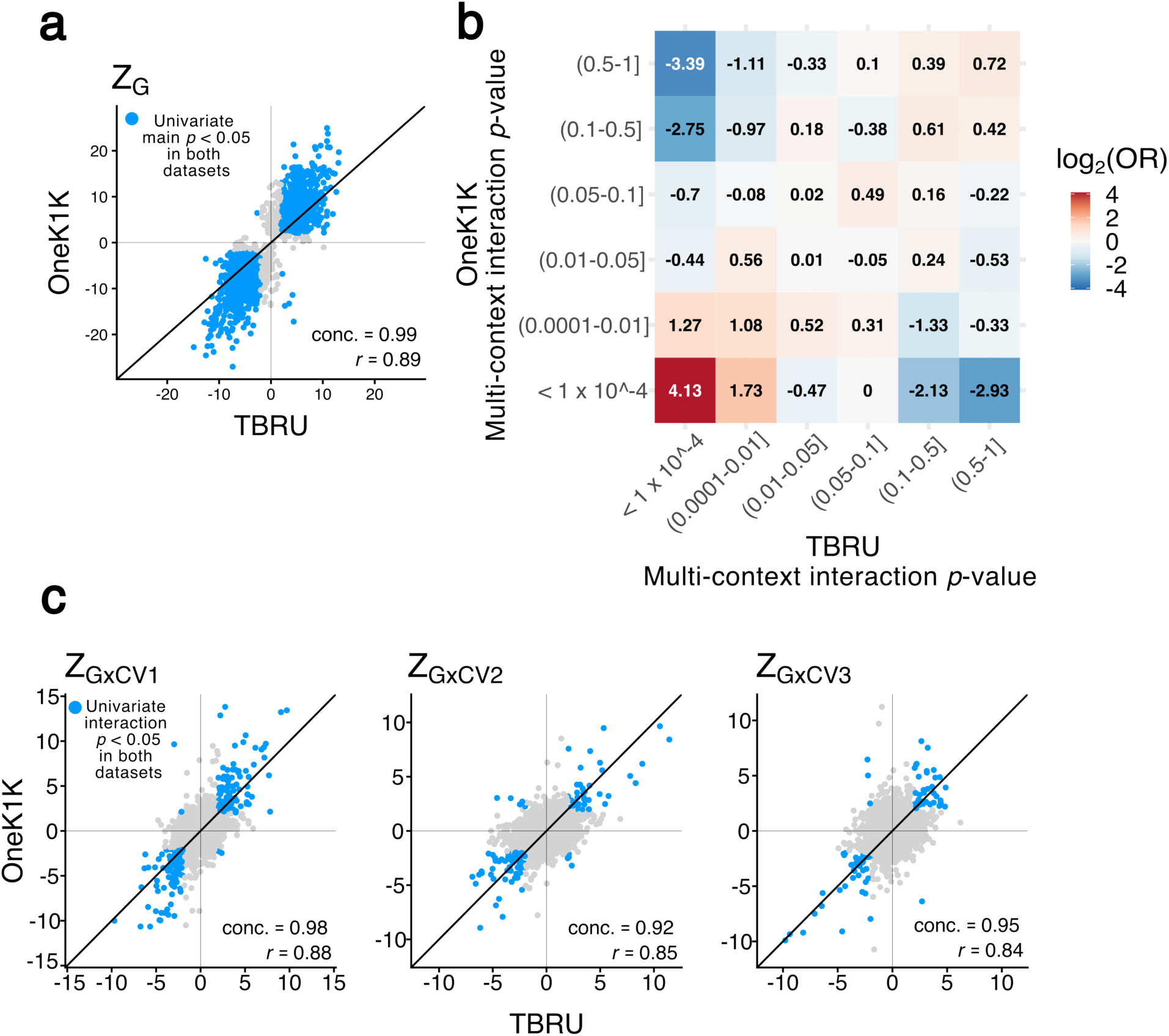
Replication of main, multi-context interaction, and single-context eQTL effects between TBRU and OneK1K with *Dynema*. **a,** Main eQTL effect *z*-scores in TBRU (x-axis) and OneK1K (y-axis). Each point represents a variant-gene pair with significant pseudobulk meta-analyzed eQTL effect after Sǐdák correction. Variant-gene pairs that were nominally significant by *Dynema* (*p* < 0.05 both datasets) are shown in blue. Proportion of nominally significant eQTLs with shared allelic direction in both datasets (concordance), and Pearson correlation are shown in the right-bottom of the plot. **b**, Replication of multi-context interaction effects. Columns and rows represent *p*-value ranges in TBRU (x-axis) and OneK1K (y-axis). Values in each tile represent a two-sample Fisher’s exact test log_2_ odds ratio between the observed and expected number of multi-context interaction eGenes in both datasets observed in the corresponding *p*-value ranges. **c,** Single-context eQTL interaction z-scores for CV1-3 (left to right, depicted as in **a**).

Next, we assessed consistency of *multi-context interactions* (**Methods; Supplementary Table 8-9**). Variant-gene pairs with low *p*-values were shared across datasets (**Figure 3b**). For example, 83% of the 117 variants with strong interaction evidence in TBRU (*p* < 1 × 10^-4^) replicated in OneK1K (*p* < 0.05). Broadly, variant-gene pairs with *p* < 1 × 10^-4^ in both datasets were highly enriched (OR > 16); while pairs with *p* < 1 × 10^-4^ in one dataset were depleted for *p* > 0.5 in the other (OR < 1/8).

We also assessed reproducibility of individual cell state interaction components. We tested *single-context interaction effects* for each CV separately for all 1,719 variant-gene pairs in both datasets (**Supplementary Table 10-15**). Strikingly, single-context interaction effects replicated (**Figure 3c**): with concordant allelic direction (0.98, 0.92, and 0.95 for CV1, CV2 and CV3) and z-score correlation (0.88, 0.85, and 0.84) for nominally significant (*p* < 0.05 in both datasets) interactions. These results demonstrate *Dynema*’s ability to identify reproducible interactions.

Consider the rs1893813 lead main variant for *ATM*, associated with Ataxia-telangiectasia^30^. Applying *Dynema* to test a *total effect* (including a *main* and *multi-context interaction*) demonstrated significance in both data sets (*p*_TBRU_= 1.7 × 10^-7^, *p*_OneK1K_= 3.3 × 10^-37^). The rs1893813-G allele showed consistent main-effect direction, though only significant in OneK1K (*β*_TBRU_ = 0.0051, *p*_TBRU_= 0.48; *β*_OneK1K_ = 0.053, *p*_OneK1K_ =8.9 × 10^-18^; see **Figure 4a**). rs1893813-G showed a negative CV1 interaction, causing its expression effect to decline and eventually become negative with increasing cytotoxicity (*β*_TBRU_ = -0.023, *p*_TBRU_= 4.9 × 10^-7^; *β*_OneK1K_ = -0.038, *p*_OneK1K_ = 3.8 × 10^-19^).

**Figure 4.**
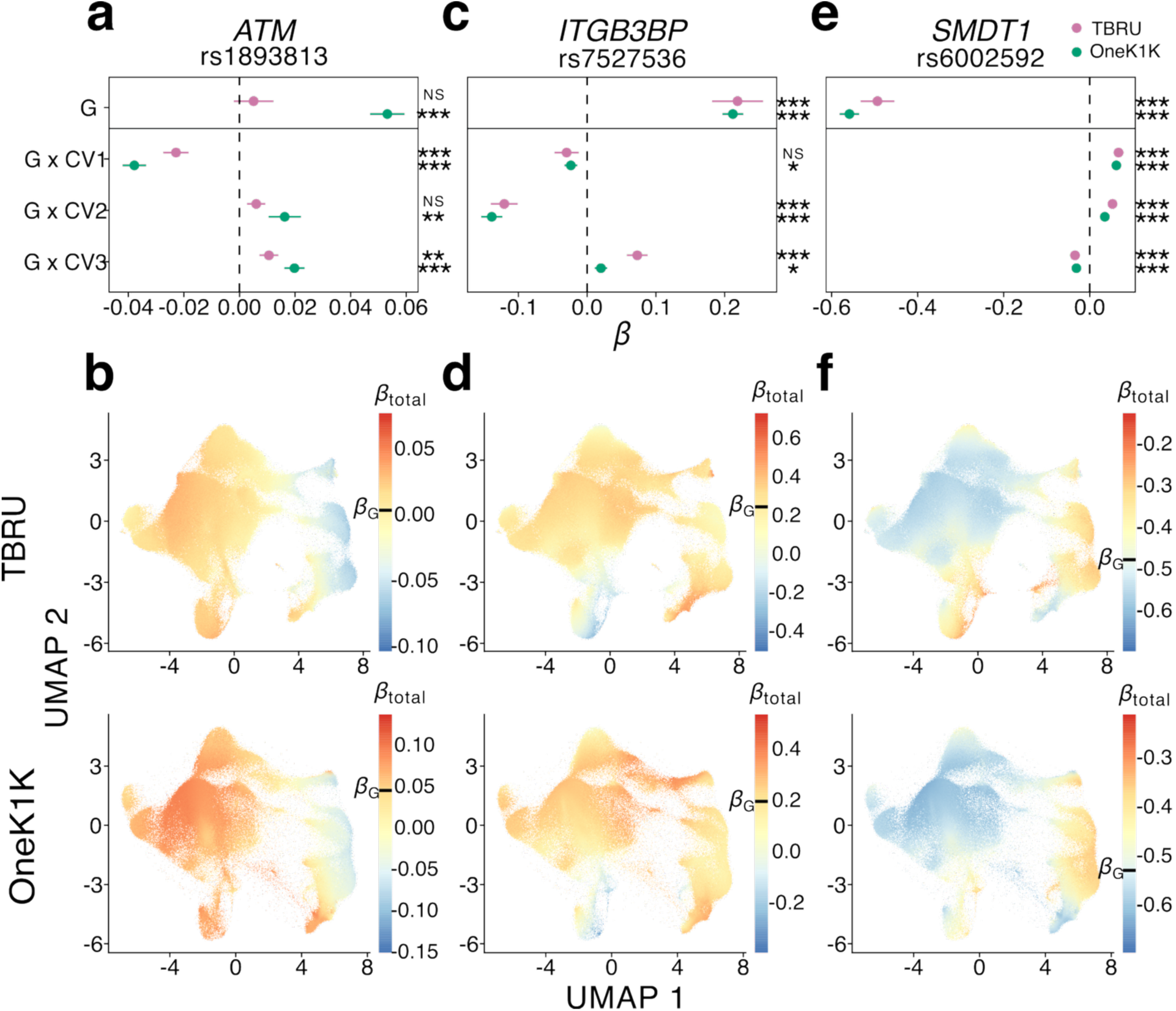
*Dynema*’s eQTL results for three highly replicable genes in TBRU and OneK1K. **a, c, e,** Forest plot of *Dynema*’s main and interaction effect sizes for CV1-3 for TBRU (pink) and OneK1K (green) for meta-analysis lead variants in *ATM*, *ITGB3BP*, and *SMDT1* loci respectively. Each dot represents the *β*value, the bars represent the 95% confidence interval, and the asterisks indicate significance of the effect according to a score/Lagrange test with *Dynema*. **b, d, f,** UMAP plots of total single-cell eQTL effect (TSCE) for *ATM*, *ITGB3BP*, and *SMDT1* respectively in TBRU (top row) and OneK1K (bottom row) datasets. Color scales are marked with a dash to indicate the main effect size, *β_G_*. Significance codes: NS (*p* ≥ 0.05), * (*p* < 0.05), ** (*p* < 0.01), *** (*p* < 0.001).

Additionally, rs1893813-G increased the eQTL effect with increasing regulatory activity (CV2) and central memory function (CV3) consistently in both datasets (**Figure 4a**). Plotting total single-cell eQTL effect (TSCE) on a UMAP reveals consistent patterns for TBRU and OneK1K (**Figure 4b**), with positive effects in central memory and helper T cells, and negative effects in cytotoxic T cells. This exemplifies an intriguing scenario where divergent effects in specific cell states are obscured by cell-state agnostic analyses in datasets where cell state proportions are balanced.

As another example, for *ITGB3BP*, rs7527536-C had its strongest cell-state interaction with CV2 in both datasets (*β*_TBRU_ = -0.12, *p*_TBRU_= 6.8 × 10^-10^; *β*_OneK1K_ = -0.14, *p*_OneK1K_ = 4.1 × 10^-19^), dampening the eQTL in regulatory T cell states (**Figure 4c-d**). At *SMDT1*, the rs6002592-C amplified the eQTL in central memory T cells (CV3) (*β*_TBRU_ = -0.034, *p*_TBRU_= 3.5 × 10^-16^; *β*_OneK1K_ = -0.031, *p*_OneK1K_ = 3.4 × 10^-20^) but significantly dampened it with increasing CV1 and CV2 (**Figure 4e-f**).

### *Dynema* identifies additional eQTLs in locus-wide cell-state-aware analysis

*Dynema*’s speed enabled single-cell resolution locus-wide eQTL analysis going beyond the lead variant. For example, for *SMDT1,* we tested total (**Figure 5a**), main (**Figure 5b**), and multi-context interaction (**Figure 5b**) effects for 949 and 1,078 variants in TBRU and OneK1K respectively (+/-250kb TSS). In this simple case, the genetic signal underlying all three tests was shared, with lead main and interaction variants in both datasets in LD with the pseudobulk meta-analyzed lead variant (rs6002592, r^2^ > 0.88) (**Figure 5d**), suggesting a common driving genetic signal.

**Figure 5.**
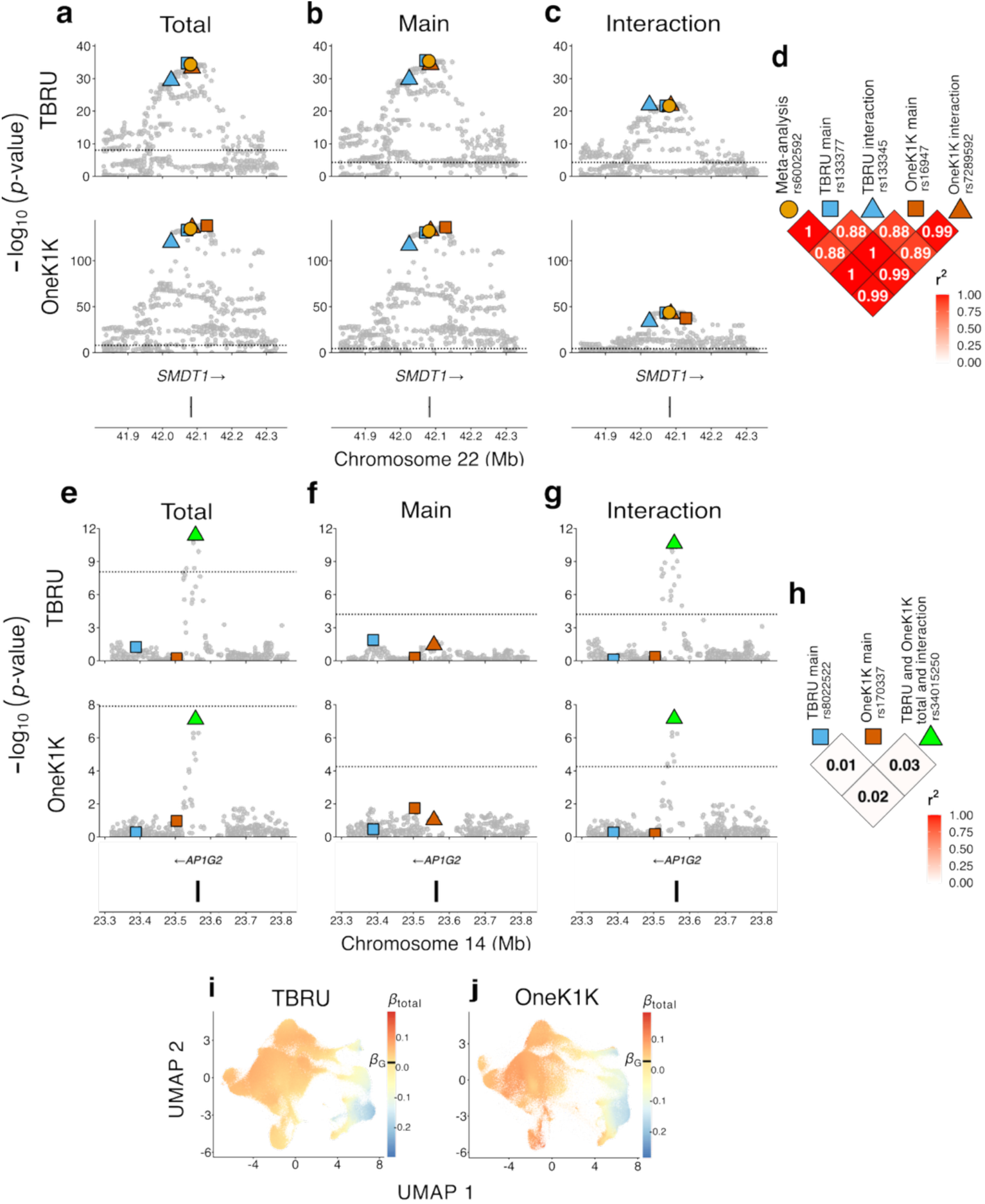
Locus-wide eQTL associations for *SMDT1* and *AP1G2* in TBRU and OneK1K. **a**, Total, **b**, main, and **c**, interaction eQTL effect locus zoom plots in TBRU (top row) and OneK1K (bottom row) for variants in the *SMDT1* locus (n_TBRU_ = 949, n_OneK1K_ = 1,078). x-axis represents the genomic coordinate, and the y-axis shows the association p-values using a score/Lagrange test with *Dynema*. Lead variants are shown for meta-analysis pseudobulk effect (yellow circle) and for the *Dynema* main (square) and interaction (triangle) effects in TBRU (blue) and OneK1K (orange). **d**, LD plot showing *r*^2^ between each pair of lead variants shown in **a**-**c**. LD information from rs73421049 was used as a proxy for rs16947 (r^2^ = 0.95 in OneK1K) due to its absence in TBRU. **e**, Total, **f**, main, and **g**, interaction eQTL effect locus zoom plots in TBRU (top row) and OneK1K (bottom row) for *AP1G2*. **h.** LD plot showing *r*^2^ between each pair of lead variants shown in **e-g. i-j,** *AP1G2* UMAP plots with total single-cell eQTL effect sizes in TBRU (**i**) and OneK1K (**j**). Color scales are marked with a dash to indicate the main effect size, *β_G_*.

We assessed if *Dynema* could uncover cell-state-dependent regulatory signals at loci without significant pseudobulk eQTL. *AP1G2* lacked a significant meta-analysis pseudobulk signal (meta-analysis p_nominal_=0.0004, *p_Šidák_*=1). But, in both datasets *Dynema* demonstrated a strong total eQTL signal for the same lead variant (rs34015250, *p*_TBRU_< 4.0 × 10^-12,^ 825 *cis*-variants tested; *p*_OneK1K_<7.5 × 10^-8^, 930 *cis*-variants, **Figure 5e**). Consistent with pseudobulk analysis, *Dynema* main effects were not significant (*p*_TBRU_>0.013, *p*_OneK1K_>0.018, **Figure 5f**). But *multi-context interaction effects*, identified the same lead variant in both datasets (rs34015250, *p*_TBRU_<2.2 × 10^-11^, *p*_OneK1K_<6.7 × 10^-8^, **Figure 5g and h**). Plotting the TSCE, we observed that the eQTL was restricted to GZMK+ T cells (**Figure 5i and j; Supplementary Figure 1**). This highlights how *Dynema* can reveal regulatory effects otherwise obscured in rare cell states.

### Lead main and interacting eQTL variants may be independent

We asked whether main and interaction eQTL effects within a locus reflect independent signals. First, we performed a genome-wide eQTL analysis to identify ‘total eGenes’ with a significant total effect correcting for variant-gene pairs tested (*p* < 8.7 × 10^-9^, 5,890,440 tests in TBRU; *p* < 1.2 × 10^-8^, 4,222,078 tests in OneK1K, **Methods**). We found 849 and 985 total eGenes in TBRU and OneK1K, respectively (**Supplementary Table 16-17**). For these, we assessed main and multi-context interaction effects, correcting for the number of variants in the *cis* window (**Methods**). In TBRU, 835 and 262 eGenes had main and interaction effects, respectively (**Figure 6a and Supplementary Table 18 and 19**). In OneK1K, eGenes had 973 and 273 main and interaction effects, respectively (**Figure 6a and Supplementary Table 20 and 21**). About 27-29% of main eGenes had a significant eQTL interaction, and ∼95% of interaction eGenes also had a main eQTL effect (**Figure 6b and c**). Intriguingly, 1-2% of total eGenes had no main effect, such as *AP1G2* (**Figure 6b and c**).

**Figure 6.**
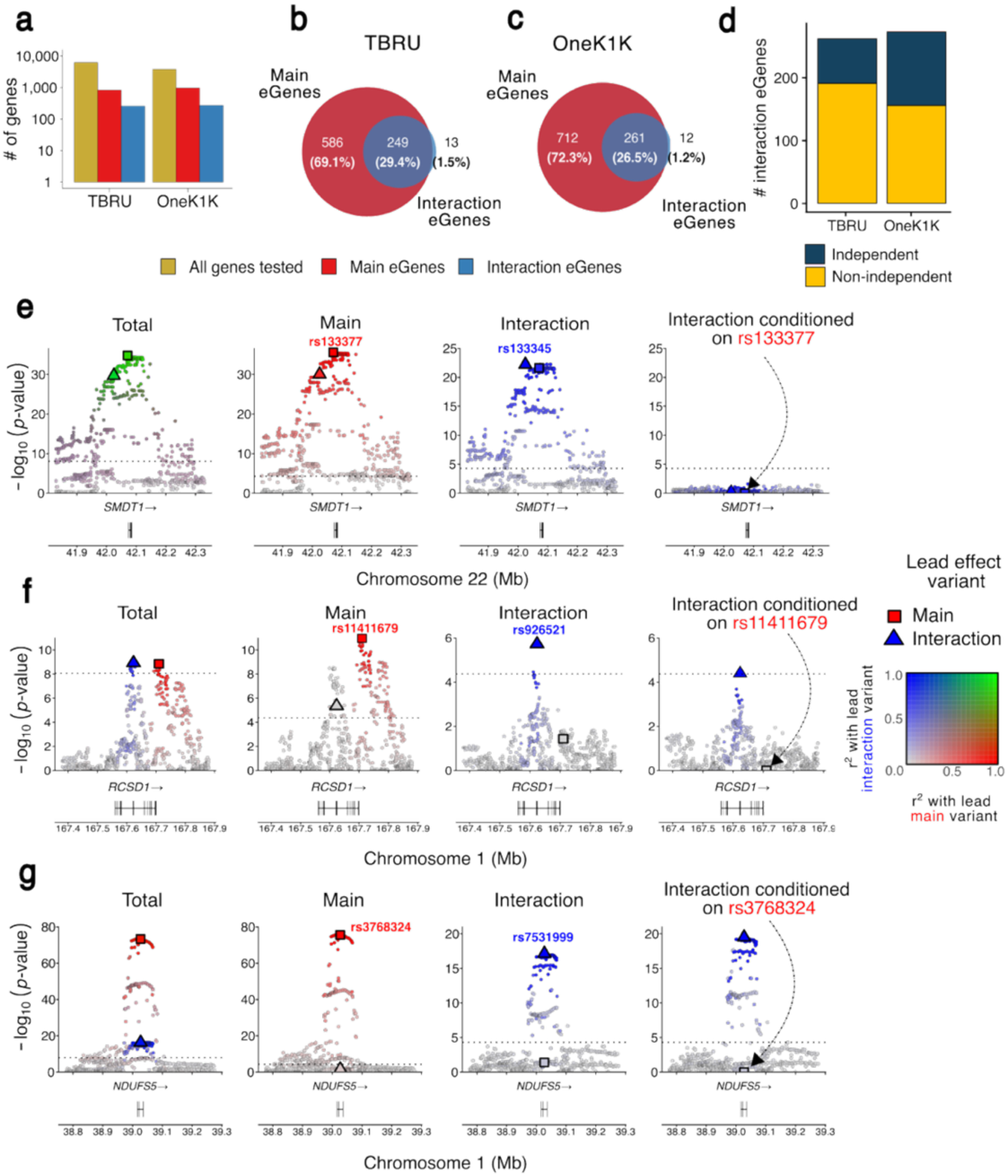
Genome-wide eQTL mapping with *Dynema* and main-effect independent interaction effects in TBRU and OneK1K. **a**, Barplot showing the total number of genes tested in each dataset in green (‘All genes tested’), genes that have at least one variant within 250kb with a significant main eQTL effect in red (‘Main eGenes’), and genes that have at least one variant within 250kb with a significant interaction eQTL effect in blue (‘Interaction eGenes’) in the TBRU and OneK1K datasets, respectively. **b**, Set overlap between main eGenes and interaction eGenes in the TBRU dataset. **c**, Set overlap between main eGenes and interaction eGenes in the OneK1K dataset. **d**, Conditionally independent cell-state-dependent eQTL effects from main effect. Each bar represents the total number of interaction eGenes detected in TBRU and OneK1K. Interaction eGenes whose interaction effects are statistically independent from the main effect are shown blue. Interaction effects explained by main effects are shown in yellow. **e-g**, Locus zoom plots of main-effect dependent interaction eQTL effect for *SMDT1* (**e**) in TBRU and main-effect independent interaction effects for *RCSD1* (**f**) in TBRU and *NDUFS5* (**g**) in OneK1K. Locus plots for total, main, interaction, and conditional interaction effects are shown for each gene. x-axis represents the genomic coordinate and the y-axis shows the association p-values using a score/Lagrange test with *Dynema*. Lead variants for main and multi-context interaction effects are shown as squares and triangles respectively. Linkage disequilibrium (r^2^) between lead and remaining variants are shown in red (main effect) and blue (interaction effect) color scales. Linkage disequilibrium between main and interaction lead variants are represented as overlay between red and blue color gradients, where high LD for both variants is represented in green.

With conditional analysis we assessed independence between interaction and main eQTL effects. We controlled for the main lead variant by including its main and CV1-3 interaction effects (**Methods**). Strikingly, 71 and 117 interaction effects remained significant in TBRU and OneK1K, respectively (*p*_TBRU_ < 1.91 × 10^-4^ ∼ 0.05 / 262 tests; *p*_OneK1K_ < 1.83 × 10^-4^ ∼ 0.05 / 273 tests; **Methods**), representing 27% and 43% of interaction eGenes (**Figure 6d**). LD between lead main and interaction variants was minimal in these loci (TBRU median r^2^ = 0.15, OneK1K median r^2^ = 0.11, **Supplementary Figure 14**). Recognizing that power to detect independence may be limited, we observed that a more permissive FDR<5% analysis suggested 50% and 58% of the interaction eQTLs were independent in TBRU and OneK1K, respectively.

We calculated the correlation between log-transformed *p*-values before and after conditioning on the main effect in each locus (**Methods**). We hypothesized that, for loci with independent main and interaction effects, conditioning should minimally affect interaction *p*-values locus-wide. As expected, median correlation of *p*-values in independent loci was high (0.87 in TBRU, 0.94 in OneK1K, **Methods; Supplementary Figure 14**).

For example, for *SMDT1* in TBRU, we observed dependence between the main and interaction effect (*p* = 5 × 10^-23^, *p*_cond_ = 0.56; **Figure 6e**); the lead variants were in high LD (r^2^ = 0.83). In contrast, *RCSD* in TBRU had a conditionally independent interaction signal (*p* = 1.8 × 10^-6^, *p*_cond_ = 3.8 × 10^-5^; **Figure 6f**), with negligible LD (r^2^ = 0.003). Similarly, *NDUFS5*’s interaction effect in OneK1K was independent of the lead main effect (*p* = 6.0 × 10^-8^, *p*_cond_ = 2.6 × 10^-20^; **Figure 6g**), with weak LD (r^2^ = 0.07). We concluded that interaction and main effects are often distinct for the same eGenes.

### Some disease variants have complex, cell-state-dependent regulatory effects

We tested colocalization of single-cell eQTL loci with seven autoimmune diseases^31–36^: Rheumatoid Arthritis (RA), Seropositive RA (RAS), Type 1 Diabetes (T1D), Systemic Lupus Erythematosus (SLE), Multiple Sclerosis (MS), Inflammatory Bowel Disease (IBD), and Grave’s Disease (GD) (**Methods).** We tested 357 TBRU and OneK1K loci that (1) had a significant total eQTL and (2) had a variant with *p_GWAS_* < 1 × 10^-7^ within the *cis* window.

We tested colocalization with total, main, and multi-context interaction eQTL effects using *Dynema*’s summary statistics (**Methods**). We identified 59 total, 60 main and 32 interaction eGenes colocalizing with an autoimmune disease locus (**Supplementary Table 22-24**). We note that about half of the colocalizations implicate a context-dependent eQTL, consistent with previous report^8^.

In some cases, the interaction effect colocalized better with disease loci than the main effect. *TSPAN32* encodes a transmembrane protein implicated in immune regulation^37^ and linked by GWAS to autoimmunity^31,38,39^ and hematological traits^37^. While expressed in hematopoietic tissues, its autoimmune mechanism is unknown^36^. The *TSPAN32* interaction effect colocalized with the seropositive RA GWAS signal (PP.H4=posterior probability of shared causal variant; PP.H4_TBRU_ = 0.99, PP.H4_OneK1K_ = 0.98), while its independent main effect did not (PP.H4_TBRU_ = 8.6 × 10^-4^, PP.H4_OneK1K_ = 1.4 × 10^-3^; **Figure 7a**); with similar results in Onek1K (**Supplementary Figure 15a**) The eQTL interaction effect was largest in the CD8+ T cells (**Figure 7b and Supplementary Figure 15b**).

**Figure 7.**
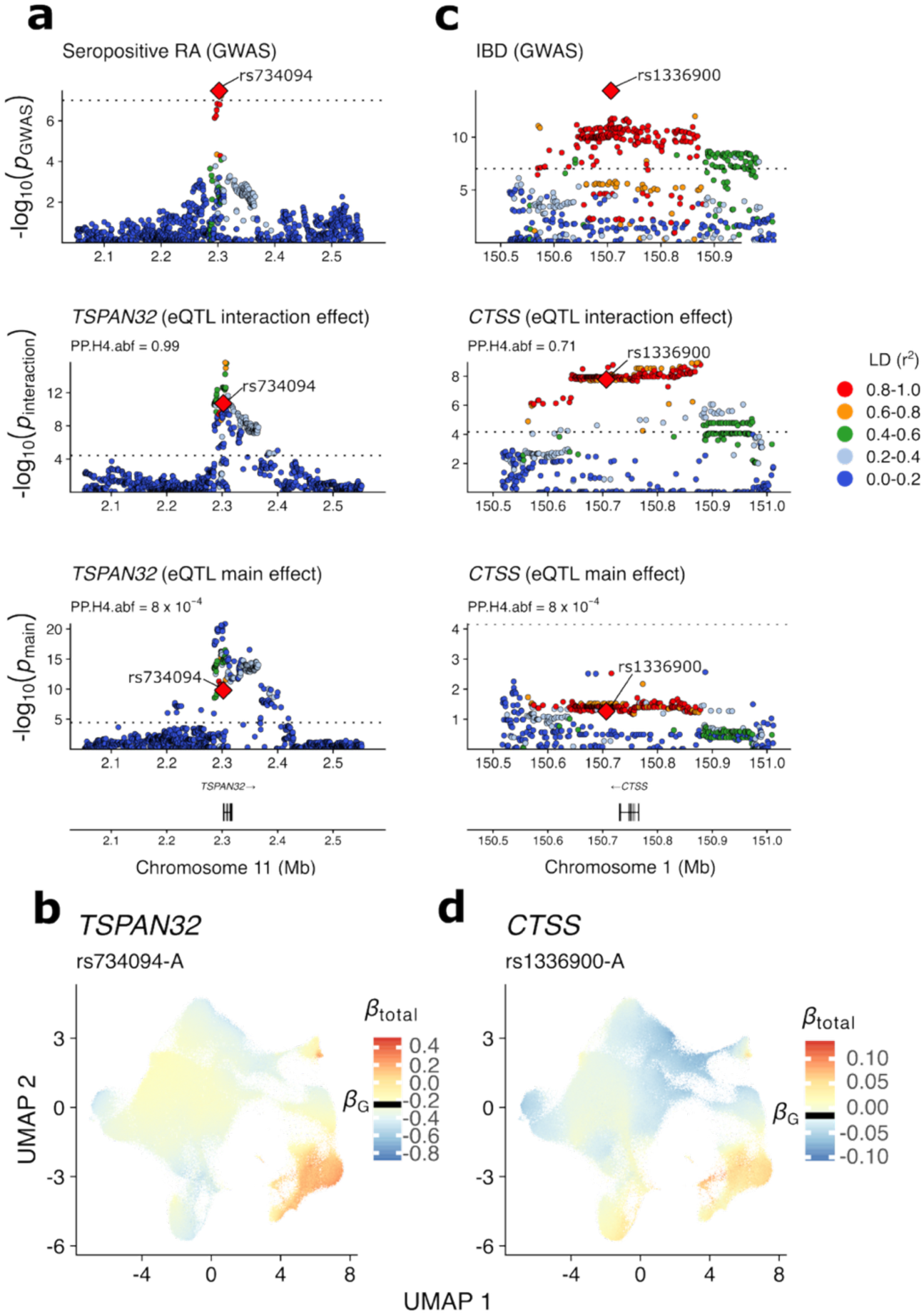
Colocalization of autoimmune disease GWAS loci with eQTL multi-context interaction effects in TBRU. Locus zoom plots are shown for autoimmune disease GWAS (top), multi-context interaction effect (middle), and main effect (bottom) in TBRU for **a,** *TSPAN32* and **c**, *CTSS* loci colocalizing with positive Rheumatoid Arthritis (RAS) and Inflammatory Bowel Disease (IBD), respectively. x-axis represents the genomic coordinate and the y-axis shows the association p-values from published GWAS or from *Dynema* for eQTL multi-context interaction and main effect tests. Each point represents a variant, and the red diamond represents the lead GWAS disease variant in each of the two loci. LD information (r^2^) of all variants with the lead GWAS is represented as color gradient. Posterior probability for common causal genetic signal between autoimmune disease and eQTL effects are shown for multi-context interaction (middle) and main (bottom) effects. **c-d**, UMAP plots with TSCE for lead interaction variants in *TSPAN32* (**b**) and *CTSS* (**d**), respectively. Color scales are marked with a dash to indicate the main effect size, *β_G_*.

*CTSS* encodes a lysosomal enzyme critical in antigen-presention^40^. Dysregulation causes chronic inflammation^40^ and autoimmunity^41–43^ . Strikingly, the eQTL interaction effect colocalized with IBD GWAS signal (PP.H4_TBRU_ = 0.71, PP.H4_OneK1K_ = 0.69); its main effect showed minimal colocalization (PP.H4_TBRU_ = 0.15, PP.H4_OneK1K_ = 2.8 × 10^-5^; **Figure 7c**). We observed similar Onek1K results (**Supplementary Figure 14c**). As with *TSPAN32*, *CTSS* exhibited independent main and interaction effects. The disease lead variant had its largest effect specifically on activated CD8+ and CD4+ T cells (**Figure 7d; Supplementary Figure 14d**), representing only ∼2% of T cells in blood.

These results highlight the potential for eQTL cell-state interaction effects to indicate potential disease allele mechanisms.

## Discussion

Here, we introduce *Dynema*, an accurate scalable method to map context-dependent eQTL effects at true single-cell resolution. We recommend users first assess the total eQTL effect at a locus to discover single-cell resolution eGenes, and then disentangle these signals into cell-state-independent (main) or state-dependent (interaction) effects. *Dynema* offers several advantages over existing methods.

First, *Dynema*’s inferential framework uses CRVEs to account for donor-level single-cell structure while preventing model misspecification; it avoids the computational costs of mixed-effects models. *Dynema* achieves genome-wide statistical calibration in contrast to other approaches and is robust to spurious cell-state-dependent eQTLs while retaining power. This is critical as disease risk alleles can have subtle, context-specific effects that are obscured in mixtures of cell states.

Second, *Dynema* is the only calibrated method fast enough for genome-wide cell-state-dependent eQTL scans. These essential analyses dissect the regulatory architecture of loci, define independent regulatory effects, and colocalize regulatory and disease associations. *Dynema*’s speed enabled identification of complex regulatory patterns discernible only with genome-wide cell-state-dependent eQTL analysis in hundreds of genes. For example, the *AP1G2* total eQTL effect is driven entirely by a cell-state-dependent regulatory signal. This eQTL effect would have been missed in pseudobulk approaches or methods that test only lead main effect variants for state-dependent effects. With conditional analyses, we show that the same locus contains independent state-dependent and state-independent effects, as in *RCSD1*.

Crucially, some disease signals are explained by context-dependent rather than main effect eQTLs. In T cells, many autoimmune disease loci colocalize with cell-state-dependent eQTLs. We identified specific examples where colocalization with SLE and RAS is driven entirely by dynamic eQTLs restricted to rare T cell subsets, rather than by independent main effects. Our data capture only resting blood T cells, potentially missing other important states associated with autoimmunity. Furthermore, we anticipate that other complex disease alleles will broadly colocalize with context-dependent eQTLs in other cell types. This could bridge the gap between eQTLs and complex disease loci by capturing effects specific to rare cell states.

Third, our study also demonstrates that context-dependent eQTLs reproduce across data sets. Demonstration of reproducibility requires accurate scalable methods, large well-powered data sets, and cell state variables that transfer between datasets. Here, for T cell data, we used canonical variates capturing transcriptional and surface marker protein signals derived in the TBRU dataset, transferring these annotations to the OneK1K dataset using reference mapping. We note that accurate definition of cell-state variables is essential for comparing and meta-analyzing single cell datasets.

*Dynema* has limitations. While CRVEs allow for an arbitrary correlation structure among cells within each donor, they assume independence between donors, preventing *Dynema*’s application to cohorts with related individuals. In practice most eQTL studies include unrelated donors, but advancing technologies will yield larger datasets where modeling relatedness may become necessary. Additionally, the robustness of CRVEs can degrade in small sample sizes^20^, and *Dynema’s* score bootstrapping^20,21^ enables calculation of accurate empirical p-values without relying on asymptotic assumptions.

We expect *Dynema* will uncover even more complex regulatory architectures in phenotypically diverse datasets, such as multi-tissue atlases. *Dynema* can capture eQTL interactions across any biological context, including cell state, perturbation responses or disease status, provided they can be represented as continuous or categorical variables at single-cell resolution. By fully leveraging this cellular resolution, researchers can better define context-dependent regulatory effects and their fundamental roles in disease risk.

## Methods

### Constructing a robust single-cell-resolution eQTL model

To accurately test associations between SNPs and single-cell gene expression, we developed a strategy based on the cluster-robust variance estimator (CRVE) and the score/Lagrange multiplier test to robustly estimate calibrated p-values in the presence of correlated cells within donors and moderate model misspecification (e.g. over- or under-dispersion). The method requires donor-level genotyping data, single-cell RNA expression data, context(s) (e.g. low-dimensional embeddings or cell weights), and donor- and cell-level covariates. Additionally, we performed the score bootstrap^15^ to refine p-value calculation for situations when asymptotic assumptions may fail.

First, we modelled single-cell gene expression counts using a Poisson Generalized Linear Model:

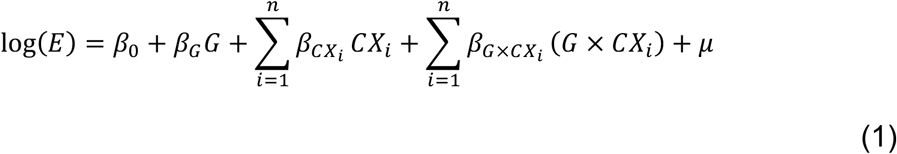

where *E* is the gene expression counts, *β*_0_is the intercept term, *G* is the alternate allele dosage of the tested variant, *CX_i_* is the context of interest, *GxCX_i_* is the interaction term between genotype and context, and *μ* is the residuals.

We added biological and technical covariates to (1) to account for possible confounding:

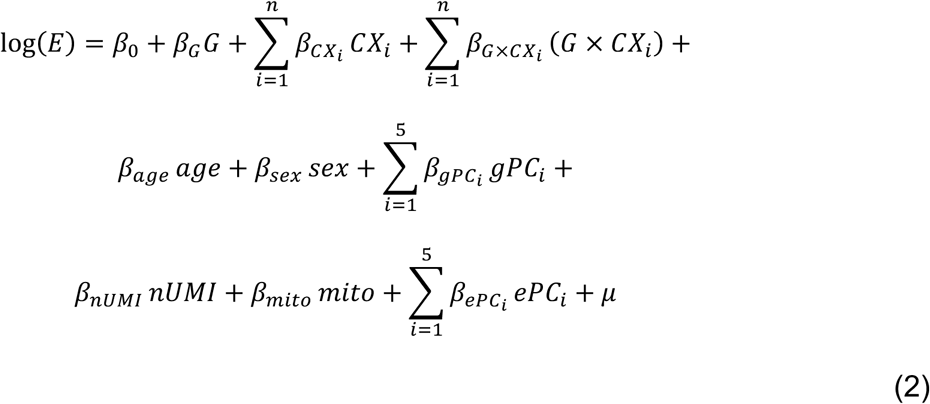

At the donor-level, *age* is the scaled age (mean = 0 and s.d. = 1), *sex* is a binary dummy variable for biological sex, and *gPC_i_* are genotype PCs to account for genetic ancestry and population stratification. At the single-cell level, *nUMI* is the scaled and natural log-transformed number of UMIs, *mito* is the proportion of mitochondrial expression, and *ePC_i_* are the expression PCs. To simplify the notation in the following steps, we can express an unrestricted (full) model (2) in its matrix form as:

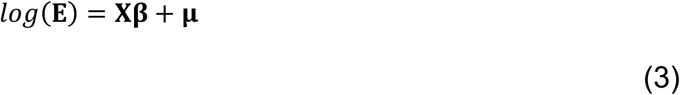

Assuming *n* cells and *p* predictor variables, *E* is an *n* × 1 column vector containing the gene expression counts, *X* is a *n* × *p* design matrix where columns each correspond to a predictor included in (2), *β* is a *p* × 1 vector, and *μ* is a column vector for the residuals.

For inferential purposes, we fitted a restricted (null) model by setting *q* terms of interest to be equal to zero:

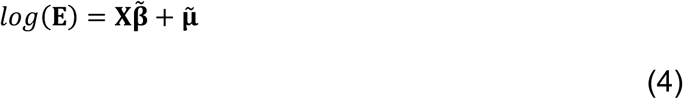

Where *β* is a vector *p* × 1 with *q* elements equal to zero, and *μ* a column vector containing the parameter coefficients and the residuals under the null model respectively.

### Computation of single-cell-level contribution to score function

To perform a Lagrange multiplier test, we first calculated the contribution of each single cell to the score function (i.e., gradient of the Poisson GLM objective log-likelihood) as proposed by Roodman *et al*. ^16^ and Kline, *et al*.^15^. The Poisson probability mass function can be defined as

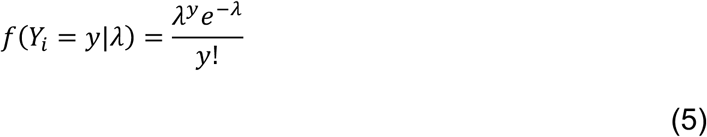

Where *λ* is the mean and variance parameter of the Poisson distribution and *y* is a discrete random variable.

The Poisson GLM log-likelihood function for each cell indexed by *i* is

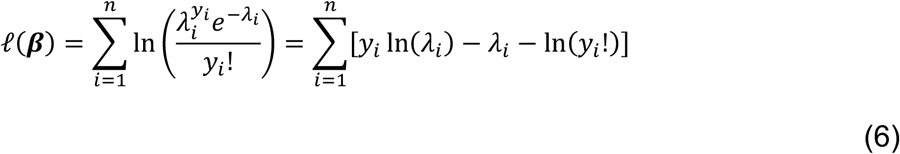

In the context of a Poisson GLM with a log-link function, we can relate *λ* to the predictors as the expected value *λ_i_* = exp(*x^T^β*); therefore, we can rewrite the log-likelihood as

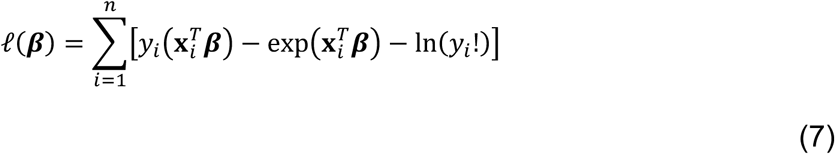

The scores *U_i_*(*β*) can be calculated by differentiating the log-likelihood function with respect to the vector *β*:

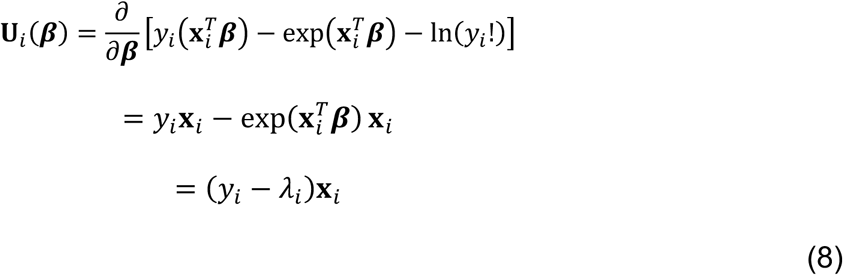

Where *y_i_* − *λ_i_* are the residuals *μ*. For parameter inference via the score test/Lagrange multiplier (LM), we built observation-level scores under the null (eq. 4) using the restricted model residuals as follows:

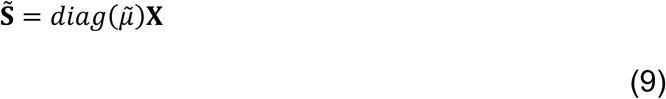

*S* is an observation-level scores matrix of dimensions *n* × *p*, where *μ* are the residuals from the restricted model. *S* contains the contribution of all predictors from the full model to the gradient of the log-likelihood function evaluated at the restricted model solution.

### Cluster-robust variance estimation

We relaxed the assumptions of the Poisson GLM to handle the correlation of cells within each donor and allow for potential model misspecification by applying the cluster-robust variance estimator (CRVE) as described in Roodman *et al.*^16^:

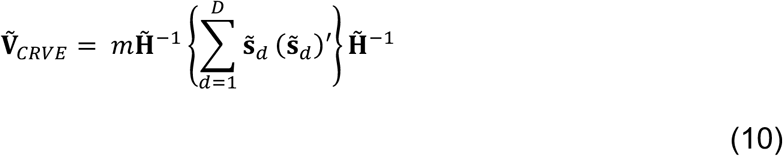

where *s̃_d_* is a vector of length *p*, the sum of all the centered single-cell scores from donor *d* for each predictor, *m* is a finite-sample correction factor *D*/(*D* − 1) (with *D* being the number of donors), and −*H̃*^−1^ is the classical Poisson GLM variance-covariance matrix of the full model evaluated at the restricted model solution:

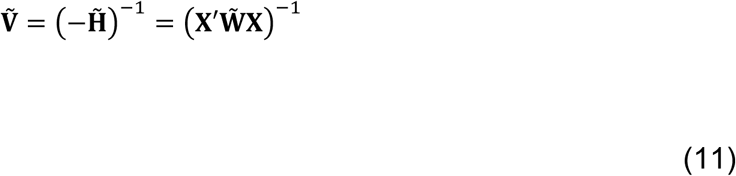

Where *X* is the design matrix with all predictors from the unrestricted model and 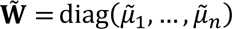 a diagonal matrix with the residuals from the restricted model.

### Lagrange multiplier / score test

We tested for single or multiple betas jointly by applying general and multiple exclusion linear restrictions. For each model, we tested the null hypothesis:

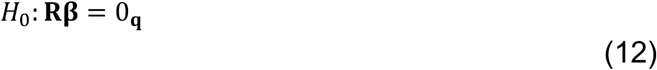

Where *R* is a matrix with *q* linear restrictions as rows and *k* parameters as columns. *R* contains 1s for the parameter(s) being tested. For example, for the model:

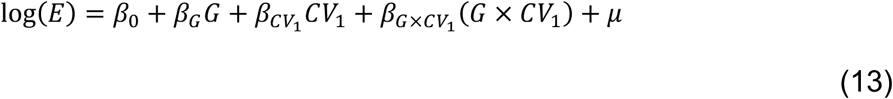

testing a single restriction (df = 1) *H*_0_: *β_G_*_×*CV*1_ = 0 requires a row matrix **R** = [0,0,0,1]. Likewise, testing the joint (df = 2) restriction *H*_0_: *β_G_* = 0, *β_G_*_×*CV*1_ = 0 requires **R** = [0,1,0,0; 0,0,0,1].

We defined the generalized Lagrange multiplier test as proposed in Wooldrige^21^:

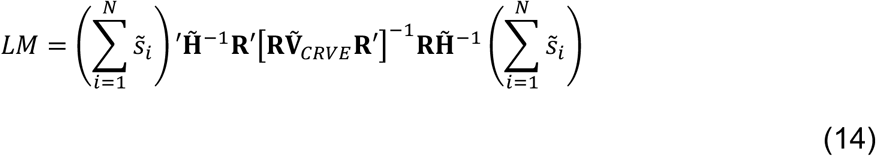

where the first and last components on the right-hand side of the formula are the sum of uncentered restricted scores for each predictor across all cells. We derived a p-value from the LM statistic based on the area under the upper tail of the *χ*^2^distribution curve with *q* degrees of freedom. When *q* = 1, the LM can be linearized to derive a Z-score.

### Adaptive score bootstrapping

When the number of individuals is small or the number of cells across individuals is considerably unbalanced, CRVEs may fail to provide accurate variance estimates^16^. For this case, we adopted as part of *Dynema* the score bootstrapping strategy proposed by Kline and Santos^15^ to derive empirical p-values without relying on analytical assumptions.

Briefly, the scores at the donor level are “perturbed” by multiplying them by random weights (either +1 or -1), which preserve the correlation structure within each donor while introducing randomness), to build an empirical distribution of the test statistics^16,15^. Since the parameters are estimated only once under the null hypothesis, this approach is faster than conventional bootstrapping and eliminates the computational burden of refitting the model for each iteration. As large *p*-values can be calculated with a smaller number of bootstrap iterations, we also implemented an adaptive procedure to reduce the computation time of the score bootstrap. We adaptively increase the number of bootstrap resamples depending on the significance level for each SNP-gene test and stop the procedure if evidence of significant association is not found.

In detail, the adaptive score bootstrapping consists of three steps:

1. For single bootstrap iteration *i*:

a. Randomly sample a “wild weight” for each donor from a Rademacher distribution {1, -1} with equal probability
b. Create a new bootstrap set of donor-level scores *S*^∗*b*^ by multiplying the scores of each donor by its corresponding wild weight
c. Calculate a bootstrap CRVE using the perturbed donor scores *S*^∗*b*^ as in eq. (10)

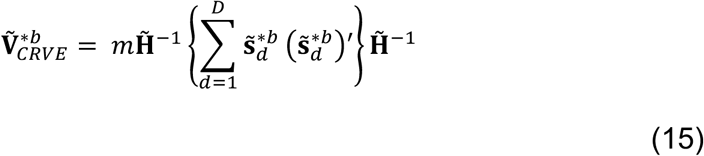
d. Compute the bootstrap *LM*^∗*b*^ test statistic using the new 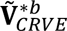 using eq. (14)
2. Repeat step 1 *b* times to create a bootstrap distribution of size *b*. If step 2 was already performed at least once, combine all *LM*^∗*b*^ computed statistics from previous rounds with the current one. Count the number of instances where *LM*^∗*b*^ statistics are equal or greater than the observed *LM*.

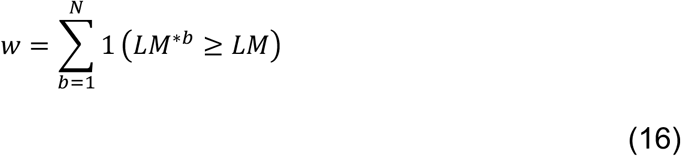
3. If *w* < 20, repeat step 2 with a greater number of bootstrap iterations *b*. If *w* ≥ 20, calculate a bootstrap p-value as follows:

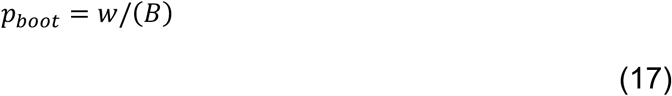

or, if the number of total bootstrap iterations is *B* + 1, as suggested in Roodman *et al*.^20^:

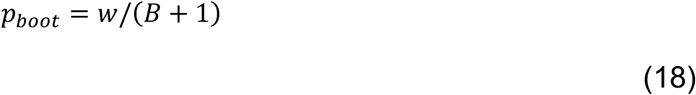

We choose a value of *w* ≥ 20 as a bootstrap stop criterion to avoid random fluctuations of the p-value near the significance level.

### Genetic data generation, processing, and quality control

#### TBRU

The data used in this study were originally generated from a Peruvian tuberculosis progression cohort. Details of the genotyping process can be found in Luo *et al*. 2019 ^20^, but in brief, 4,002 admixed Peruvian individuals were genotyped on a custom Affymetrix array (LIMAArray) designed based on whole exome sequencing from 116 individuals from the same Peruvian cohort.

We used the processed genotyping data from Nathan, *et al*. 2022^8^, which included 675,358 genotyped variants after removing variants significantly associated with batch (p < 1 × 10^-5^), duplicated, with low call rate (< 95%), significant differences in the missingness rate between cases and controls (> 10^-5^), or Hardy-Weinberg p value < 10^-5^ in controls. These 675,358 variants were mapped to GRCh37/b37, pre-phased with SHAPEIT2^44^, and imputed with IMPUTE2^45^ using the 1000 Genomes Project Phase 3 as the reference panel. After removing variants with an INFO score < .9, minor allele frequency < 0.05, or deletions, the remaining variants were converted to GRCh38 with liftOver^46^ , resulting in 5,486,956 remaining variants for eQTL analysis.

Genotyping principal components were previously calculated as described in Nathan, *et al*. 2022^8^, by combining autosomal genotyping data from the full cohort (n = 4,002) with whole-genome sequencing data from the 1000 Genomes Project Phase 3 (n=2,045 individuals from 26 populations); matching variants with MAF > 1% on position, reference, and alternate allele; removing highly correlated variants (r^2^ > 0.2, window size = 50, and step size = 5); and conducting PCA with PLINK^47^.

#### OneK1K

The data used in this study was originally generated as part of the OneK1K project. Details of the genotyping process can be found in Yazar *et al*. 2022^12^. Briefly, 1,034 individuals were genotyped using the Illumina Infinium Global Screening Array.

We used the processed genotyping data from Rumker *et al*. 2024^48^, which included 492,431 genotyped variants after removing duplicated variants, with low call rate (≤98%), or Hardy-Weinberg *p*-value < 10^-6^. These 492,431 variants were mapped to GRCh37/b37, pre-phased with SHAPEIT2^44^, and imputed using Minimac3^49^using the 1000 Genomes Project Phase 3 as the reference panel. After removing variants with an INFO score < .7, minor allele frequency < 0.05, the remaining variants were converted to GRCh38 with liftOver^46^ , resulting in 4,227,088 remaining variants for eQTL analysis.

Genotyping principal components were previously calculated as described in Rumker *et al*. 2024^48^, by combining autosomal genotyping data with whole-genome sequencing data from the 1000 Genomes Project Phase 3 (n=2,045 individuals from 26 populations); conducting PCA with PLINK^47^. After intersecting the genotype data with the single-cell RNA-seq dataset and removing ancestry outliers based on genetic PCs as described in Yazar *et al*. 2022^12^ (individuals ± 6 SD away from the European mean of PC1 and PC2), we retained 969 individuals.

### Single-cell RNA and CITE-seq data generation, processing, and quality control

#### TBRU

The memory T cell CITE-seq dataset used in this study was previously generated as part of a study of a subset of the genotyped Peruvian tuberculosis progression cohort ^21^. All samples were collected from individuals after treatment and resolution of *M. tuberculosis* infection. Detailed experimental methods are available in Nathan, *et al*. 2021. In summary, we collected PBMCs from 259 genotyped Peruvian individuals, isolated memory T cells through negative selection on CD45RA, and carried out an optimized version of the CITE-seq protocol^50^ with a panel of 31 surface proteins with TotalSeq^TM^-A (BioLegend) oligonucleotide-labeled antibodies. We conducted 10x Genomics library preparation and sequencing for pools of six donors and subsequently demultiplexed donors using Demuxlet^51^.

In the present study, we used single-cell sequencing data from Nathan, *et al*. 2021^8^ that was aligned to GRCh38 with Cell Ranger (3.1.0). Antibody tags were aligned to a dictionary of oligonucleotides. We followed the same quality control steps as in the original study: we removed doublets labeled by Demuxlet^51^ and other genotype mismatches, cells with <500 genes expressed or >20% of mitochondrial unique molecular identifiers, and cells lacking memory T cell surface markers (CD3 and CD45RO). After QC, 500,089 cells remained. We log-normalized gene expression data and applied centered log-ratio transformation to surface marker UMI counts for each of these cells.

We used canonical variates (CVs) reflecting an RNA- and protein-informed low-dimensional embedding of cells that were published in Nathan, *et al*. 2021 to represent cell states. These CVs were defined through canonical correlation analysis^52^ (CCA) on single-cell expression of genes (union of the top 1,000 most variable genes per sequencing batch, excluding TCR genes) and non-control surface markers. We computed 2-D UMAP embeddings from the top 20 CVs using the uwot R package^53^ with n_neighbors = 30L, metric = “euclidean”, min_dist = .1.

#### OneK1K

The single-cell RNA-seq dataset used in this study was previously generated as part of the OneK1K project. Detailed experimental methods are available in Yazar *et al*. 2022^12^. In this study, we used the pre-processed single-cell RNA-seq from Rumker *et al*. 2024^48^. In summary, PBMCs were collected for 969 genotyped Australian individuals. Individuals were processed across 75 10x Genomics libraries, sequenced. Single-cell transcriptomes were subsequently demultiplexed using Demuxlet^51^. Single-cell sequencing data was aligned to GRCh38 with Cell Ranger (2.2.0). We removed cells with< 660 UMIs, ≥ 7.8% mitochondrial reads, and ≤ 230 genes detected. We stringently removed doublets following recommendations in Kang *et al*. 2023^11^. Briefly, we considered doublets called by either Demuxlet^51^ or Scrublet^54^. We retained CD4+ and CD8+ T cells (including the naïve, memory, and effector compartments) based on the original labels reported the OneK1K authors. After QC, ∼579,000 cells remained. We log-normalized and scaled (mean = 0 and standard deviation = 1) gene expression data

UMI counts for each of these cells. We determined 3,000 highly variable genes a performed PCA analysis using Seurat^55^. We selected the top five principal components as covariates for eQTL analysis.

### Defining TBRU’s continuous cell states in OneK1K with reference mapping

We defined T cell states in the OneK1K dataset by transferring cell state information (CV1-3) from the TBRU dataset using Symphony^26^. First, we built a T cell Symphony reference using TBRU’s CCA output including embeddings, loadings, mean and standard deviation of log-normalized expression data for each gene, and batch information (i.e. donor). Symphony creates harmonized reference CV embeddings (hCVs) that can be used to map a query dataset. For data visualization, we also stored 2-D UMAP embeddings into the reference object to also enable UMAP projection of query datasets.

We mapped OneK1K T cells to the TBRU Symphony reference using the *mapQuery* function, providing OneK1K’s gene expression data and donor information. We subset the first three reference-mapped CVs (CV1-3) and used them as the T cell states for eQTL analysis in OneK1K. Additionally, we generated projection coordinates for the OneK1K dataset in the TBRU reference UMAP embedding using Symphony.

### Assesing *Dynema*’s calibration for context-dependent eQTL testing

We designed two simulation strategies to create null datasets from a set of 2,202 pseudobulk eQTLs identified in Nathan *et al.*^8^ : 1) permutation of genotypes across donors and 2) simulation of gene expression data under a null parametric model:

1) *Genotype permutation.* We randomly shuffled genotypes across donors and expanded them at the single-cell level. This strategy disrupts genetic effects for main and interaction terms. All remaining covariates are kept unchanged.
2) *Gene expression simulation.* We simulated gene expression data under the null (no cell-state-dependent eQTL interaction) for a variant-gene pair as follows:

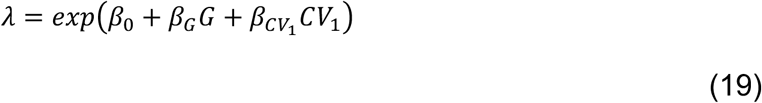

Where *β*_0_ = *log*(*θ* + 1) (with *θ* being the observed average gene expression for each gene) and *G* is the alternate allele dosage for the variant. We randomly sample UMI counts from a Poisson distribution based on the simulated *λ*.

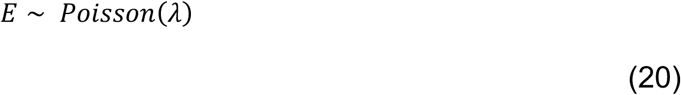

We simulated gene expression under four null scenarios where *β_GxCV_*_1_ = 0 and calculated analytical and bootstrap *p*-values with *Dynema*:

1. No main effect, no differential expression. *β_G_* = 0 and *β_CV_*_1_ = 0
2. No main effect, differential expression. *β_G_* = 0 and *β_CV_*_1_ = 0.2
3. Main effect, no differential expression. *β_G_* = 0.2 and *β_CV_*_1_ = 0
4. Main effect, differential expression. *β_G_* = 0.2 and *β_CV_*_1_ = 0.2

We also evaluated *Dynema*’s calibration when multiple contexts were modeled by testing:

1. *Single-cell state interaction*: *GxCV*1 with simulated expression data
2. *Multi-cell state interaction*: *GxCV*1, *GxCV*2, and *GxCV*3 using permuted genotypes and simulated expression data
3. *Main genetic effect*: G and no interaction terms modeled using permuted genotypes.
4. *Total genetic effects*: *G*, *GxCV*1, *GxCV*2, and *GxCV*3 using permuted genotypes

We calculated the observed analytical and bootstrap *p*-values with *Dynema* and created QQ-plots to compare them to the expected *p*-values from a uniform null distribution. We calculated lambda inflation factors (*λ*) to quantify inflation by transforming the observed analytical and bootstrap p-values to a *χ*^2^ statistics with n-degrees of freedom (i.e. df = 1, df = 3, or df = 4 depending on the test) using the inverse cumulative distribution function and dividing the median of the observed *χ*^2^ statistics by expected median of null *χ*^2^statistics.

### Benchmarking *Dynema*’s calibration and speed benchmarking compared to other eQTL models

#### Null: *β*G = 0 and *β*CV1 = 0

We compared *Dynema*’s calibration and speed with other existing approaches for detecting context-dependent eQTL interactions by testing for G x CV1 interactions using the permuted-genotypes null dataset. For each null scenario, we computed a false positive rate as the proportion of significant *p*-values (*p* < 0.01) and a lambda inflation factor by obtaining the median of all observed χ^2^ statistics (calculated from the observed *p*-values with *q* degrees of freedom according to each test) divided by the theoretical median of a χ^2^ distribution with *q* degrees of freedom. We calculated the total computation time for each test using a single CPU core using a HPC cluster.

Each model was implemented as described below:

#### CASTIE

We fitted a null model including donor-level covariates (age, biological sex, and five genetic PCs), and single-cell covariates (% of mitochondrial expression, and five expression PCs, and CV1). We used log(nUMIs) as an offset as recommended by the authors^27^. We specified CV1 as a single-cell context covariate. We built a LD-genome-wide panel of 3,000 markers with PLINK2^47^ pruned (--indep-pairwise 50 5 0.2) to estimate null-model variance components. We tested G x CV1 interaction using a score test. We restricted CASTIE’s analyses to 2,122 as CASTIE failed computation for 80 variant-gene pairs with non-alphanumeric gene names (e.g. *TRAV8-1* or *ZNF529-AS1*).

#### *scPME* (Single-cell Poisson Mixed-Effect model)

We applied our scPME model previously proposed in Nathan *et al*. ^8^ . We modelled UMI counts including the covariates in (2) and using a random intercept effect for donor as follows:

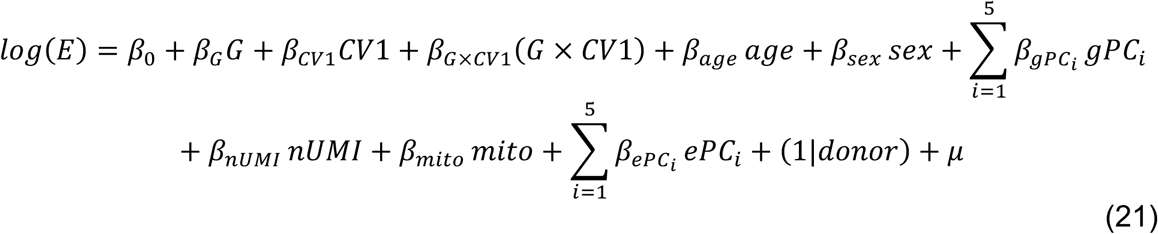

To test the significance of the eQTL interaction, we fitted a null model without the interaction term:

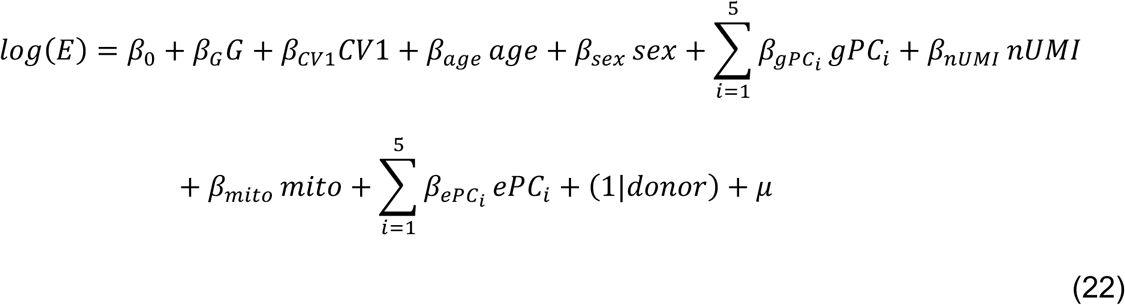

We derived a *p*-value using a likelihood ratio test between the full and null model. The scPME model was applied using the MixedModels.jl^56^ library in Julia^57^.

#### *scLME* (Single-cell Linear Mixed-Effect model)

We normalized UMI counts by dividing raw counts by each cell’s library size and multiplying by a size factor of 10,000. We added a pseudocount of 1 and applied a natural log-transformation. We modeled UMI counts using a mixed-effect model with a random intercept effect for donor as follows:

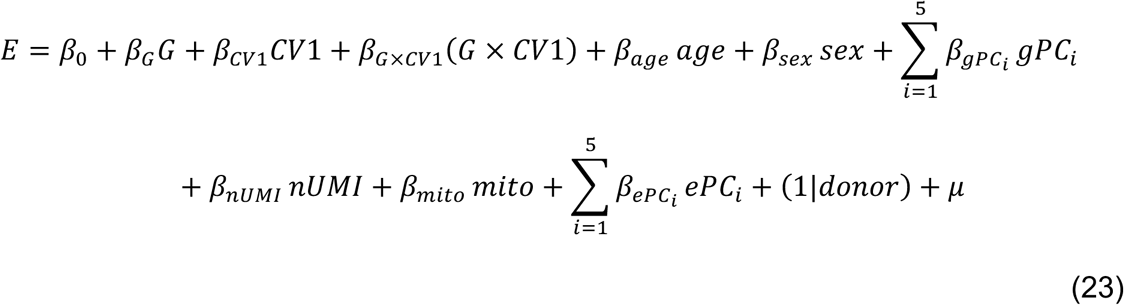

Similarly, we fitted a null model without the interaction term:

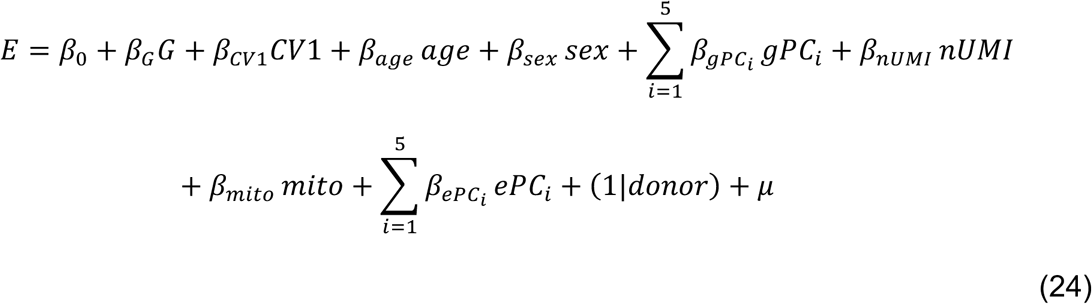

We computed p-values using a likelihood ratio test between both models. The scLME model was applied using the MixedModels.jl library in Julia^57^.

#### CellRegMap

We normalized gene expression following the same strategy used for the scLME model. We ran *CellRegMap* using the same covariates as in eq. (2) and modelled CV1 as a random effect. We tested for GxCV1 interaction using a score test as described in Cuomo *et al*., 2022.

Null: *β*G > 0 and *β*CV1 = 0 and *β*G > 0 and *β*CV1-3 = 0

We extended our parametric Poisson model-based simulation (eq. 19; **Supplementary Figure 5 and 7**) to assess the calibration of interaction effect tests in the presence of weak and strong main effects by varying *β*_G_ values under realistic effect sizes from 0 to 0.5 in increments of 0.1. We used the same set of 2,202 variant-gene pairs and simulated gene expression data for each pair as follows:

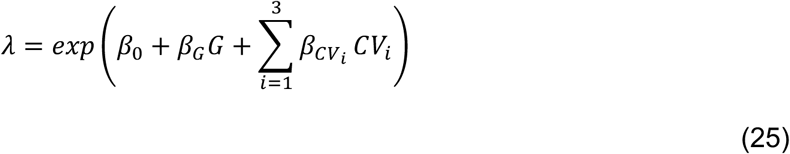

Where β_0_ = log(θ + 1), with θ being the observed average expression of each gene, and G is the alternate allele dosage of the tested variant. For the multi-context interaction test, we set all cell-state differential expression terms to β_CVi_ = 0.2. For the single-context interaction test for CV1, we used eq. 19 and β_CV1_ = 0.2. No genotype-by-context interaction term is included in this model, so the ground-truth interaction effect is exactly zero for every gene at every value of β_G_. UMI counts were sampled from a Poisson distribution parameterized by the simulated λ (eq. 20).

As a complementary data-driven approach, we simulated main effects via binomial thinning of real count data as suggested by Gerard (2020)^28^ . We used the same 2,202 variant-gene pairs and the same main-effect dose grid used in the parametric simulation (β_G_ ∈ {0, 0.1, 0.2, 0.3, 0.4, 0.5}). We used the set of permuted genotypes used for the remaining calibration analysis prior to thinning. Because the permutation procedure is independent of all other covariates and of any unmeasured biological or technical sources of variation, it removes any genotype-expression association present in the real data while preserving the data’s true biological and technical variance structure. For each gene, we downsampled UMI molecules from each cell’s observed count using a retention probability that depended only on the donor’s permuted genotype dosage at the tested variant. We defined retention probabilities for each genotype by centering the linear predictor, subtracting the maximum genotype effect of the allele dosage value (G_i_ = 2) as suggested by Gerard (2020)^28^ :

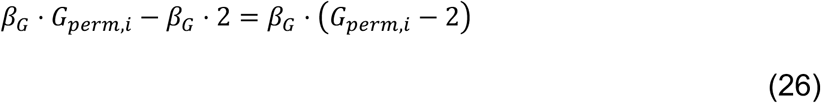

With retention probability defined as:

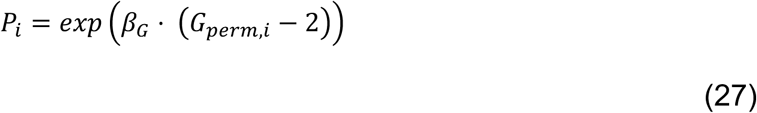

where G_perm,i_ is the permuted alternate allele dosage of the donor of cell i. Donors homozygous for the alternate allele (G_perm,i_ = 2) retain all UMI molecules at the maximum probability P_i_ = 1, heterozygous donors (G_perm,i_ = 1) retain molecules at P_i_ = exp(-β_G_), and donors homozygous for the reference allele (G_i_ = 0) at the minimum probability P_i_ = exp(-2β_G_). Retention probability decays as allele dosage decreases. For each gene, each of a cell’s real observed UMI molecules c_i_ was independently retained with probability P_i_:

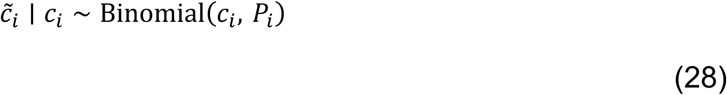

Where c̃_i_ are the resulting thinned UMI counts with simulated genotype main effect of size β_G_.

We used *Dynema* to test multi-context interactions with CV1-3 using a score test with df = 3 across all simulation approaches, and null β_GxCV1-3_ = 0 scenarios where β_G_ ≥ 0.

For calibration comparison, we applied CASTIE to test the same null single-context (G x CV1) and multi-context (G x CV1-3) interactions across the same simulation two approaches. We estimated CASTIE’s null-model variance components using an LD-pruned genome-wide panel of markers rather than the full genotype set, to avoid proximal contamination between the variance-component estimate and the variant being tested, following the authors’ recommendation^27^ . This panel was built once from the full imputed TBRU genotype data (autosomes only, MAF ≥ 0.05, dosages hard-called at a 0.49999 threshold, duplicate variant IDs collapsed to their first record) by LD-pruning with PLINK2^47^ (--indep-pairwise 50 5 0.2: 50-variant windows, 5-variant step, pairwise r^2^ < 0.2) and then randomly subsampling the pruned set to 3,000 markers (seed 42). We followed the two-step procedure from CASTIE’s tutorial for each variant-gene test. In step 1, we included CV1, and CV1, CV2, and CV3 as cell-level contexts via –dynamicCovarColList (for single-contexts and multi-context tests, respectively), used --traitType=count, and provided the pruned marker set via --plinkFile. For consistency, we used the same argument values for the remaining parameters as suggested by the authors. For the parametric Poisson simulated data, we included biological sex as sample-level covariate via --sampleCovarColList as CASTIE returned an error if no sample-level covariates. Likewise, we included sex a covariate for *Dynema* for exact comparison. For the binomial thinning simulation, we used the same sample-level covariates as for *Dynema* (sex, age, five genetic PCs, five expression

PCs, and CV-13) and included the log(total nUMIs) per cell as an offset term (as suggested by the authors). In step 2, for the parametric Poisson simulated data, we provided the original genotypes (no permutation) for each of the 2,202 variant-gene pairs. For the binomial thinning simulation, we provided the same set of permuted genotypes used throughout the benchmarking calibration analyses. Because CASTIE requires hard-called allele dosages from PLINK binary genotype files, we set -- minMAF=0 and --minMAC=0.5 to ensure all variants were tested after converting imputed dosages. With the default thresholds (--minMAF=0.05, --minMAC=5), a variant could otherwise be dropped if its MAF or MAC fell slightly below these cutoffs after hard-calling. We restricted CASTIE’s analyses to 2,122 as CASTIE failed computation for 80 variant-gene pairs with non-alphanumeric gene names (e.g. *TRAV8-1* or *ZNF529-AS1*).

We quantified inflation by computing a genomic inflation factor (λ) across the range of simulated main-effect sizes. To compute λ, we used degrees of freedom appropriate to each method’s interaction test. For *Dynema*, we used df = 3 for multi-context interactions and df = 1 for single-context interactions. For CASTIE, we used df = 1 for all interactions, as its multi-context interaction test combines per-context p-values via the aggregated Cauchy combination test (ACAT-O) into a variant-level p-value (pval_ge_CCT column in output). We created QQ plots to compare the observed distribution of interaction *p*-values to the expected uniform null for each method.

Expected *p*-values and λ calculations were performed independently for each method to account for the difference in the number of tested variant-gene pairs (n = 2,202 for *Dynema*, and n = 2,122 for CASTIE).

### Assessing calibration in downsampling conditions

We created five downsampled datasets using our genotype-permuted null dataset. First, we downsampled to 200, 100, or 50 donors. Next, we considered all 259 donors and in only a half of them, we downsampled cells by selecting a random subset of 10%, 25%, or 50% of cells per donor. Finally, we created a dataset by downsampling to only 50 donors and in only a half of them, we downsampled cells by selecting a random subset of 10%, 25%, or 50% of cells per donor. We applied *Dynema* in each of these datasets and computed both analytical and bootstrap *p*-values. We quantified statistical inflation in all scenarios by computing lambda inflation factor (λ) as described before.

### Calculating power to detect context-dependent eQTLs

We evaluated *Dynema*’s statistical power to detect GxCV interactions by simulating gene expression data across a range of realistic values for three genes (*MAF*, *FCGR3A*, *BLK*) representing high, medium, and low expression, with average number of UMI counts *θ* = 0.345, 0.054, and 0.004 respectively. We simulated gene expression as follows:

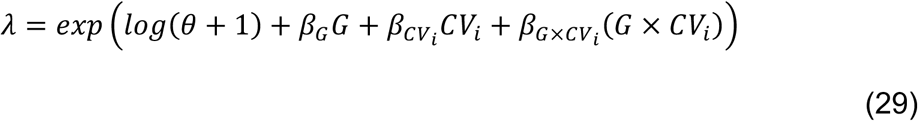

where *θ* is the baseline average UMI count expression, *G* is the allele dosage of variant with minor allele frequency = 0.2, and *CV*1 is the cytotoxic T cell state in TBRU. For all simulations, we fixed *β_G_*= 0.2 and *β_CV_*_1_= 0.2, and evaluated power at a range of *β_GxCV_*_1_ values from 0 to 0.01 with step increments of 0.0025. For a given instance of *β_GxCV_*_1_, we randomly sample from a Poisson distribution with *λ* = *λ_sim_* to simulate UMI counts.

Next, we used *Dynema* to estimate CV1-dependent eQTL effects in the simulated gene expression data. We repeated these steps 1000 times and quantified power by counting the number of instances where *p* < *α*, with *α*= 0.05 ,0.001, and 1 × 10^-4^.

### Identifying cell-state interactions for pseudobulk lead variants

We created pseudobulk gene expression profiles for each donor in each of the TBRU and OneK1K datasets, respectively. For each donor, we summed up the total number of UMIs across all their T cells for each gene. We restricted our analysis to autosomal genes with at least one UMI detected in at least 5% of T cells and 50% of the donors in each of the datasets, respectively. We normalized gene expression for systematic differences in library size across donors by computing CPM (counts per million) values and applied a log_2_ transformation. To ensure that the expression profiles across donors were normally distributed, we applied an inverse normal transformation (INT) across each gene. We residualized gene expression with respect to the top five genetic PCs, age, biological sex, and up to 45 PEER^58^ factors. We performed pseudobulk eQTL mapping with FastQTL^59^(v2.184) across all genes for variants within +/- 250Kb *cis* region from the transcription start site (TSS) of each gene in each of the datasets separately. We tested 6,289 and 3,821 expressed genes in TBRU and OneK1K, respectively. We nominated lead variants for each gene based on the smallest nominal p-value observed within the *cis* window. We applied a strict multiple hypothesis correction across the total number of variant-gene pairs using a Šídák correction^29^, *p*_Sidak_ = 1 − (1 − *p*)*^m^*, where *m* is total number of variant-gene pairs tested in each dataset (n_TBRU_ = 5,890,440 and n_OneK1K_ = 4,227,088 variant-gene pairs). We applied a significance level threshold of 0.05 on adjusted *p*-values to call significant variant-gene pairs.

We applied *Dynema* to test each variant-gene pair for multi-context interaction effects (df = 3) with cell states CV1-3 as follows:

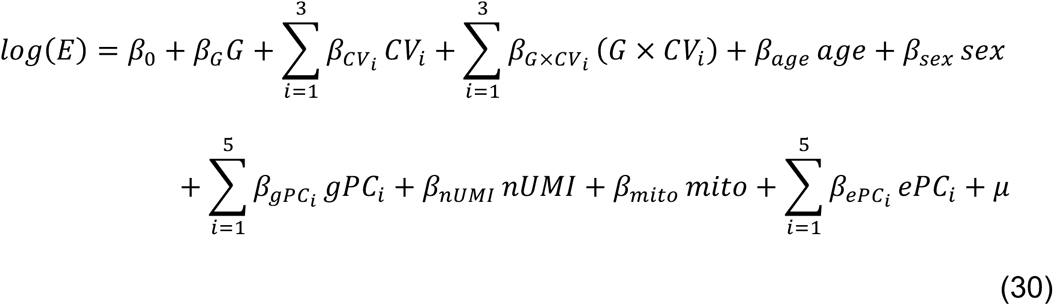

We computed Storey *q*-values^60^ for each of the lead variants and identified significant interaction eGenes with *q* < 0.05.

### Pseudobulk eQTL meta-analysis of TBRU and OneK1K

To obtain a robust set of lead variants, we performed a pseudobulk eQTL meta-analysis combining the TBRU and OneK1K FastQTL summary statistics computed previously.

We restricted meta-analysis to the intersection of 3,740 (at least one UMI detected in at least 5% of T cells and 50% of the donors in each of the datasets, respectively). We used the inverse-variance-weighted average approach^61^ to compute meta-analyzed betas, standard errors, and z-scores as follows:

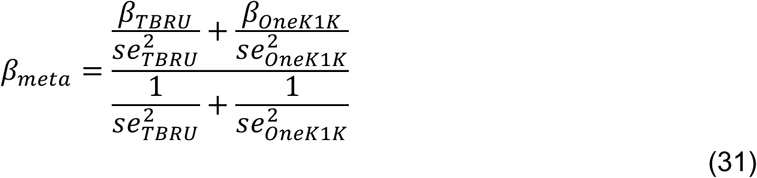

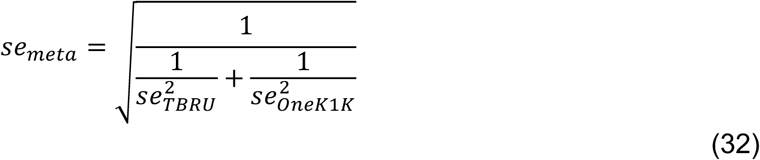

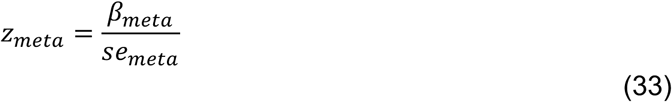

We identified lead variants for each gene based on the smallest *p*-value. We called significant meta-analyzed lead variants using a Bonferroni-Šídák correction (*p* < 1.7 × 10-8; n = 2,939,514 variant-gene pairs shared in both datasets).

### Assessing reproducibility of eQTL interaction effects across datasets

We applied *Dynema* to test the pseudobulk meta-analysis eQTLs for main (df = 1) effects (no interaction terms) as follows:

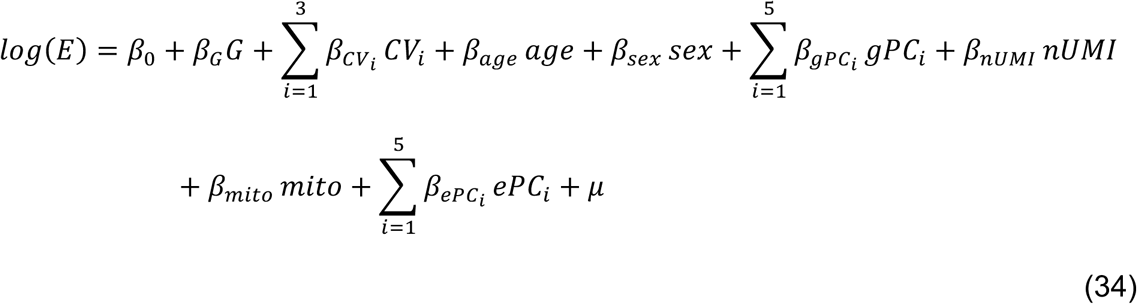

We tested multi-context interactions (df = 3) using eq. 30. Additionally, we tested independent single-context interactions with each of the three cell states (k = 1, 2, or 3) as follows:

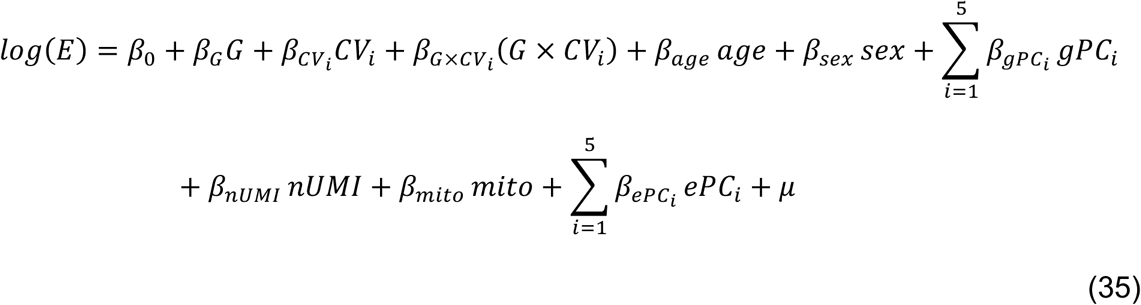

For single-context interaction tests, we quantified two reproducibility metrics: (1) concordance of allelic directionality, i.e., proportion of eQTLs with same sign for the *β* of interest in TBRU and OneK1K, and (2) Pearson correlation between *β* values from the two datasets across all pseudobulk meta-analysis lead variants with *p*_Dynema_ < 0.05 (nominally significant) in both datasets.

For the multi-context test (df=3), we also quantified replication of eGenes in both datasets by quantifying the overlap in eQTLs across different *p*-value ranges (0.5 – 1, 0.1 – 0.5, 0.05 – 0.01, 0.01– 0.05, 0.0001–0.01, < 1 × 10^-4^) using a contingency 2×2 table. For each pairwise comparison of p-value ranges between TBRU and OneK1K, we calculated a log _2_ odds ratio (log₂OR) a from a two-sample Fisher’s exact test using Maximum Likelihood Estimation conditional on the fixed margins of the table.

### Identifying main and interaction eGenes with *Dynema* with genome-wide analyses

We applied *Dynema* to identify ‘total eGenes’ with any eQTL effect using a total eQTL effect test (df=4) across all *cis* variants for each gene in each of the TBRU and OneK1K datasets using eq. 2 testing for the joint effect of the main and CV1-3 interaction components. We adjusted nominal *p*-values by applying a genome-wide Sǐdák correction across all variant-gene pairs (n_TBRU_ = 5,890,440 and n_OneK1K_ = 4,227,088).

We defined ‘total eGenes’ as genes where the lead variant had an adjusted *p*-value < 0.05. Next, we tested main (df=1) and multi-context interaction (df=3) effects with *Dynema* for all total eGenes in both datasets. We applied a locus-wide Sǐdák correction for all *cis variants* (m = total number of variants <250kb from the gene’s TSS). For each test, we defined ‘main’ or ‘interaction eGenes’ if the locus-wide adjusted *p* < 0.05, respectively.

### Conditional eQTL mapping with *Dynema*

We performed conditional eQTL mapping to determine whether an interaction eQTL effect was independent from the main eQTL effect for a given gene. We tested all *cis* variants, for each gene (+/-250kb from the TSS) for an interaction eQTL effect while conditioning on the main and interaction effects of the gene’s lead main eQTL variant as follows:

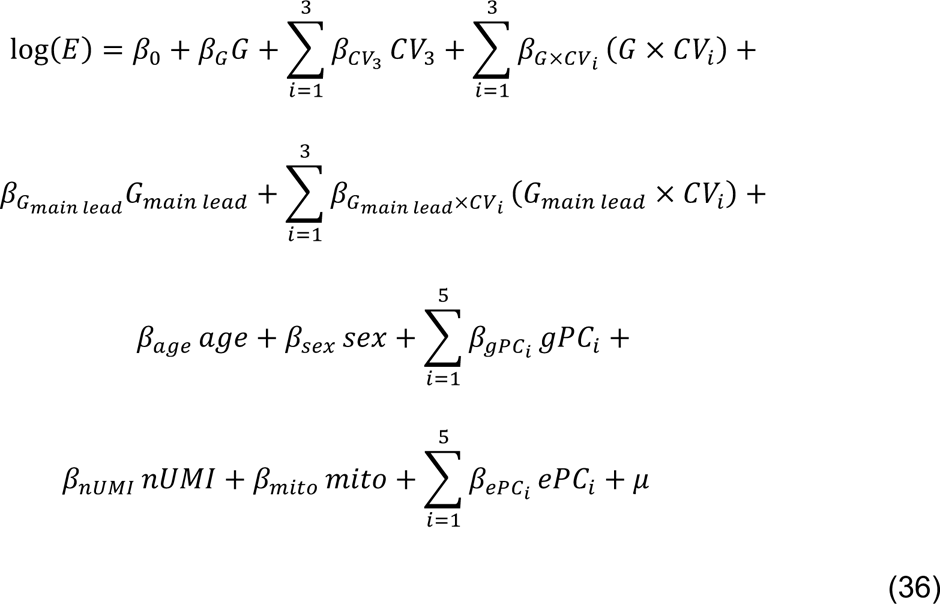

Where *G* is the tested cis-variant and G_main lead_ is the main effect lead variant previously identified using a main effect interaction test for the same gene. We computed conditional multi-context interaction effect (df=3) *p*-values for each variant. We defined a gene as having conditionally independent interaction effects if the conditional eQTL mapping results met two criteria: 1) the locus-wide Bonferroni-corrected p-value for the original lead interaction effect variant remained < 0.05 after conditioning, and 2) the correlation between log-transformed interaction effect p-values and log-transformed conditional interaction effect p-values across variants was > 0.9. For *RCSD1* and *NDUFS5*, we calculated the LD r^2^ between the lead main and lead interaction variant in TBRU and OneK1K respectively.

### Colocalization with autoimmune disease GWAS

We obtained GWAS summary statistics for seven autoimmune diseases from published studies: Rheumatoid Arthritis^31^ (RA), Rheumatoid Arthritis Seropositive^31^ (RAS), Type 1 Diabetes^32^ (T1D), Systemic Lupus Erythematosus^33^ (SLE), Multiple Sclerosis^34^ (MS), Inflammatory Bowel Disease^35^ (IBD), and Grave’s Disease^36^ (GD). We restricted our colocalization analysis to loci with a GWAS lead variant with *p* < 1 × 10^-7^ within the +/- 250Kb *cis* window. After intersecting all ‘total eGenes’ and GWAs loci, we performed colocalization on a set of 357 GWAS loci.

Then, we used coloc^59^ (version 5.2.3) to test colocalization of total, main, and interaction eQTL effects for all ‘total eGenes’ identified with *Dynema* in both TBRU (849) and OneK1K (935) with loci associated with these seven autoimmune diseases. We define colocalization events as those with a posterior probability of H4 (“both traits are associated and share a single causal variant”) > 0.5 in at least one of the TBRU or OneK1K datasets. We created locus plots using the locuszoomr^62^ package. We plotted the TSCE for *TSPAN32* and *CTSS* as described before.

### Computational implementation of *Dynema*

*Dynema* is implemented as library in the Julia programming language^57^ and can be installed from https://github.com/immunogenomics/Dynema.jl. The adaptive score bootstrapping used in *Dynema* is adapted from the WildBootTests.jl library^63^.

### Code availability

Code for the analyses conducted in this work can be found in https://github.com/immunogenomics/Dynema_analysis.

## Supplementary notes

### (1) *Dynema* is well-calibrated to test interaction, main, and total effects under different null scenarios

We assessed *Dynema*’s calibration to test single-context interactions under four additional null scenarios defined by the presence or absence of the main genetic and/or differential gene expression effect across cell state (*i.e*. β_G_ = 0 and/or β_CV1_ = 0). To capture the diversity of genes, we simulated gene expression for each of the 2,202 variant-gene pairs using observed mean expression and genotype information (**Methods**). Both analytical and bootstrapped p-values from *Dynema* showed robust calibration across all null scenarios (0.93 ≤ λ ≤ 1.04; **Supplementary Figure 5**).

We then validated *Dynema*’s statistical calibration to test more complex regulatory effects by testing multi-context interaction effects. First, we repeated our initial analysis of permuted genotypes while including interaction terms in our model for CV1, T Regulatory/Activation (CV2) and Central Memory (CV3) states. We tested multi-context interaction effects and again observed well-calibrated p-values (λ = 1.01; **Supplementary Figure 6**). We also observed well-calibrated p-values for multi-context interaction effects under the other null scenarios with simulated main and/or CV effects but no interactions (1.02 ≤ λ ≤ 1.05; **Supplementary Figure 7**). Finally, we evaluated *Dynema*’s calibration to test only main and total effects. We applied *Dynema* to our genotype-permuted set of eQTLs and observed robust calibration across all genes tested for main effects alone and total effects (λ = 1.09 and λ = 1.06, respectively; **Supplementary Figure 8 and 9**). These results demonstrate that *Dynema* is statistically calibrated to test main, single and multi-context interaction, and total eQTL effects under different null scenarios.

### (2) *Dynema*’s eQTL interaction statistics are well-calibrated in the presence of main effect eQTLs

A well-calibrated context-dependent single-cell eQTL mapping method must accurately distinguish between main (context-independent) and interaction (context-dependent) effects. Main effect eQTLs are common throughout the genome, and most do not interact with cell states^8,64,65^. Statistical models may inadvertently attribute main effects to an interaction effect, particularly for genes that are differentially expressed between contexts but have no context-dependent genetic drivers of the gene’s expression (Nathan *et al.* 2022^8^). The distinction between main and interaction effects is crucial to accurately nominate causal biological contexts (*e.g*. cell states) in which disease variants have regulatory effects.

Therefore, for methods that demonstrated sufficient calibration of context-interaction *p*-values when there was no main effect (*i.e*., β_G_ = 0), we assessed the calibration of interaction *p*-values in the presence of main effects of varying magnitudes. We built null multi-context-interaction scenarios (*i.e*., *β*_G_ ≥ 0 and β_GxCV1-3_ = 0) using two complementary strategies:

1. a fully parametric Poisson simulation (**Supplementary Figure 5 and 7; Methods**). We generated synthetic single-cell counts from a Poisson model with realistic main genotype effects (*β_G_*) and no interaction term, with no biological or technical noise assumptions (**Methods**).
2. a data resampling approach, *i.e*. binomial thinning^28^ . We leveraged real single-cell count data from TBRU, removed any genetic effect (main and interaction) via genotype permutation, and then selectively downsampled UMIs according to each donor’s permuted genotype. Distinct downsample probabilities per genotype were defined as a function of the target main effect that we simulated (**Methods**). This procedure induces a main effect in the absence of interaction effects while leaving the data’s actual biological and technical noise structure intact, providing an additional layer of evidence to complement the simplifying assumptions of parametric simulation. We used the same set of 2,202 variant-gene pairs leveraged throughout the benchmarking analyses, and we quantified calibration by calculating the percent of significant tests at *p* < 0.01 (*i.e*., for a well-calibrated method, we expect that only 1% of interaction-test *p*-values should be significant in these null interaction scenarios).

We simulated six scenarios using both simulation methods: no main effect *β*_G_ = 0 and a range of five different realistic main effects (*β*_G_ = 0.1, 0.2, 0.3, 0.4, 0.5). We observed that *Dynema*’s interaction *p*-values remain well-calibrated across all main effect sizes using both simulation approaches across null scenarios with *β*_G_ = 0 or *β*_G_ > 0 (**Supplementary Figure 10**). This was true when we tested interactions with all three CVs in aggregate (**Supplementary Figure 10a-b**) and when we tested for interactions with CV1 alone (**Figure 2d-e**); these results were consistent with our previous calibration analysis (**Supplementary Figure 5**). The percent of tests with *p* < 0.01 ranged from 0.64% to 1.14% across scenarios and simulation approaches (**Supplementary Figure 10b**). We also computed a genomic inflation factor (λ), ranging from 1.01 to 1.07 (analytical) and from 0.96 to 1.03 (bootstrap) across all null *β*_G_ ≠ 0 scenarios for both simulation approaches (**Supplementary Figure 10a**).

We also tested CASTIE^27^ (Liu *et al.* 2026, *medRxiv*). CASTIE also demonstrated adequate calibration for interaction effects in the absence of main effects (**Figure 2a**). We further evaluated its calibration under this more challenging null scenario with non-zero main effects. Under the null scenario with *β*_GXCV1-3_ = 0 and *β*_G_= 0, we recapitulated the calibration results reported in the original manuscript^27^: *p*-values are calibrated or moderately deflated (λ=0.73 and 0.96 for parametric Poisson and binomial thinning simulations, respectively; **Supplementary Figure 10a**). However, CASTIE detects a substantial proportion of false positives under the null scenarios where *β*_G_ > 0. The interaction *p*-values become increasingly inflated as the main effect grows stronger in both simulation approaches, indicating that CASTIE’s test severely and systematically conflates genuine main effects with interaction signals. This is true when testing interactions with CV1 (**Figure 2d-e**) alone and when testing CV1-3 interactions in aggregate (**Supplementary Figure 10a**). Across all null *β*_G_ > 0 scenarios and simulation approaches, λ values ranged from 1.12 to 20.35 and the proportion of significant tests for the presence of an interaction at *p* < 0.01 ranged from 6.41% (for very small main effects *β*_G_ = 0.1) to >70% (with larger but reasonably sized main effects in the Poisson simulation; **Supplementary Figure 10b**).

For each method, we observed concordant results with both simulation approaches, which suggests that each method’s calibration results reflect genuine performance rather than artifacts of either simulation’s specific assumptions. These simulation strategies were necessary because the null scenario (*β*_GxCV1-3_ = 0 and *β*_G_ > 0) cannot be directly assessed via genotype permutation alone, which ablates both the main and interaction effects simultaneously.

These results highlight the importance of evaluating calibration of context-dependent single-cell eQTL models under multiple null models that reflect realistic biological scenarios.

### (3) *Dynema* results are generally consistent with analysis with single cell poisson mixed effects model

We evaluated *Dynema*’s ability to recapitulate main and interaction eQTL effects reported in our previous TBRU eQTL study^8^. First, we assessed *Dynema*’s ability to identify main eQTL effects for the 6,511 eQTLs identified via pseudobulk analysis in Nathan *et al*. These eQTLs should have an average effect across all cells, without accounting for any cell-state dependent effects. We compared the main effect eQTL *p*-values and effect sizes between *Dynema* and FastQTL pseudobulk and observed strong concordance between these approaches (**Supplementary Figure 12a**), recapitulating 96% of main eQTL effects (nominal *p* < 0.05 in both datasets). We confirmed that *Dynema*’s main effects were concordant with the results of the scPME model (**Supplementary Figure 12b**), recapitulating 98% of main eQTL effects (nominal *p* < 0.05 in both datasets). We observed that z-score statistics from scPME were systematically higher than from *Dynema* (Deming regression *β* = 1.26). While this may indicate that *Dynema* has reduced power relative to scPME, it may also reflect *Dynema*’s better statistical calibration due to its robustness to misspecification.

Next, we used *Dynema* to test single-context interaction with CV1 for these 6,511 eQTLs. We compared *Dynema*’s GxCV1 interaction effects with those estimated by the scPME model. We observed that z-scores were correlated (*r* = 0.88) but were approximately half the magnitude of z-scores from scPME (Deming regression *β* = 1.97) (**Supplementary Figure 12c**) recapitulating 66% of interactions significant with scPME (*p* < 0.05 in both datasets). The reduced z scores for GxCV1 interactions by *Dynema* may reflect better p-value calibration, but also some loss of power which is expected with a model that is more robust to model misspecification (**Supplementary Figure 12c**). Nevertheless, the concordance of allelic direction and Pearson correlation between both methods were high (100% and 0.88 respectively) for the significant eQTLs (*p* < 0.05 for both methods) demonstrating concordant interaction signals.

