## Supplementary Figures for "Efficient genome-wide mapping of reproducible, context-dependent eQTLs at single-cell resolution"

### Supplementary information

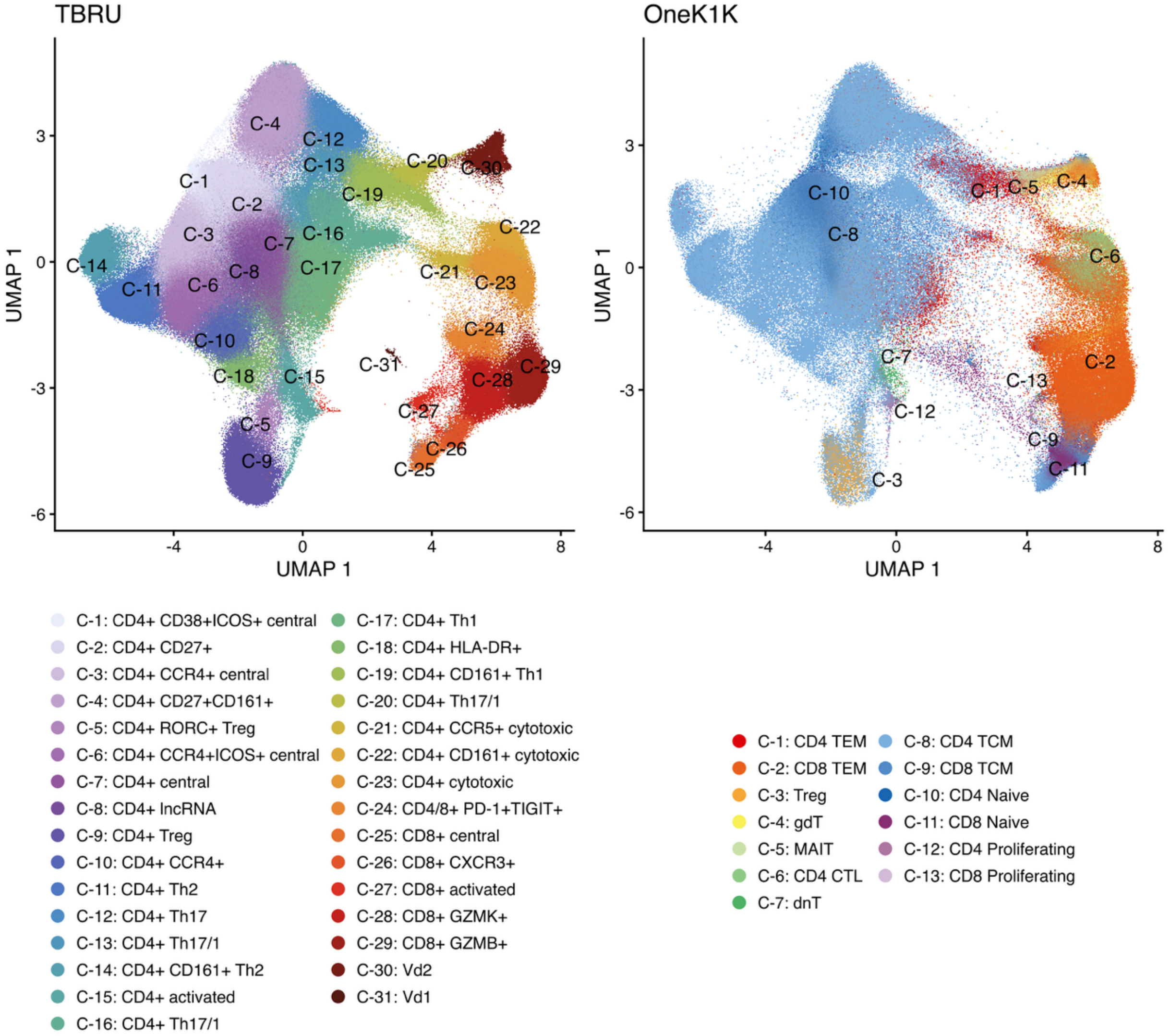


**Supplementary Figure 1 | Cell state clusters in TBRU and OneK1K**. UMAPs colored by cluster labels from original publications for TBRU (left) and OneK1K (right).

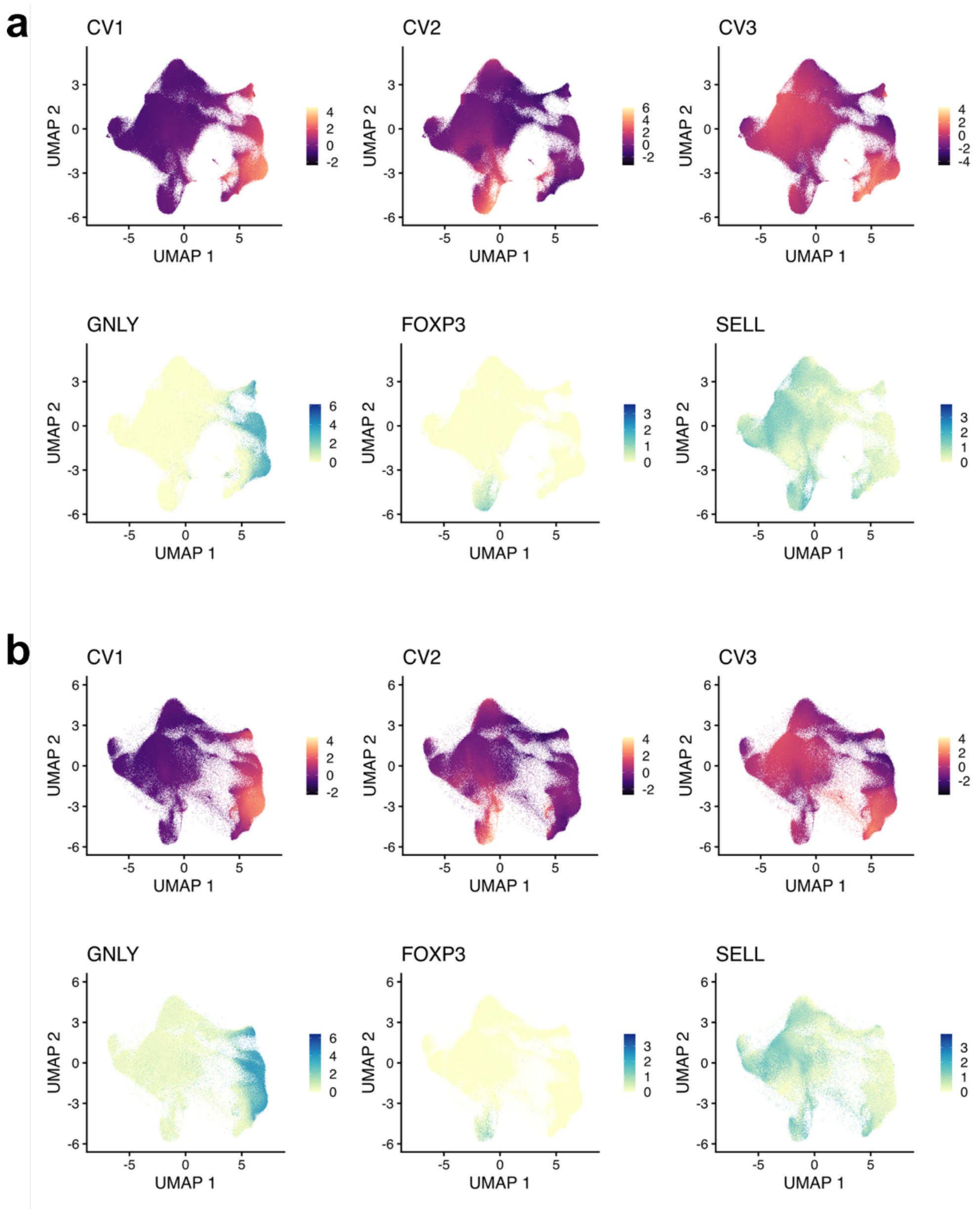


**Supplementary Figure 2 | Cell state variables ($CV_{1-3}$) and expression of gene markers in TBRU and Onek1K.** a) UMAPs colored by CVs corresponding to cytotoxicity (CV1), regulatory T cell and/or activation (CV2), and central memory (CV3) phenotypes in the TBRU dataset. Gene markers distinguishing each cell state are also shown. b) Same as above for the OneK1K dataset, where CVs 1-3 were inferred using Symphony with the TBRU dataset as reference.

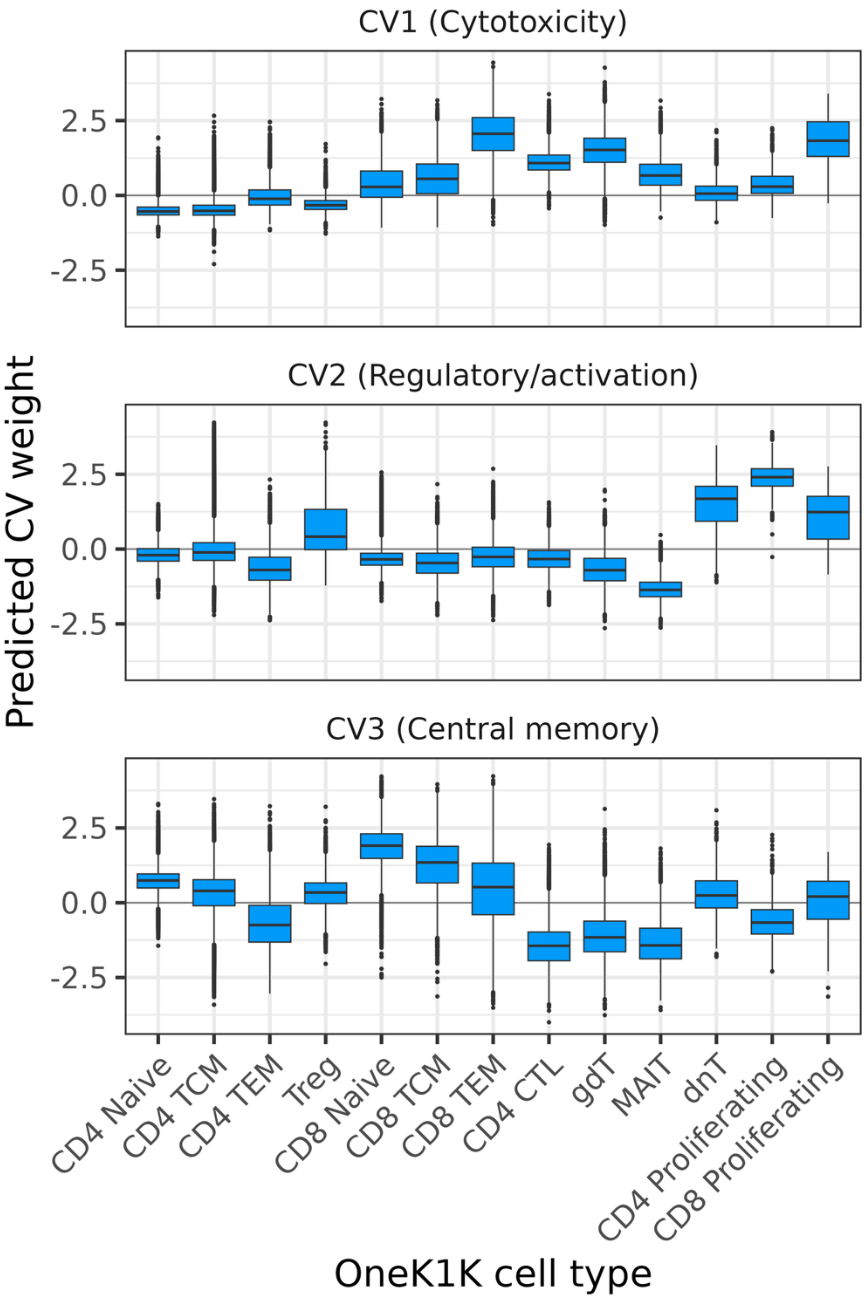


**Supplementary Figure 3 | Predicted cell state (CV) in OneK1K T cell clusters**. Each panel corresponds to a cell state (CV1, CV2, CV3) predicted in OneK1K T cells by using *Symphony* with TBRU as reference. The boxplots show the distribution of the cell state variable stratified by cluster labels from OneK1K reported by the authors (x axis).

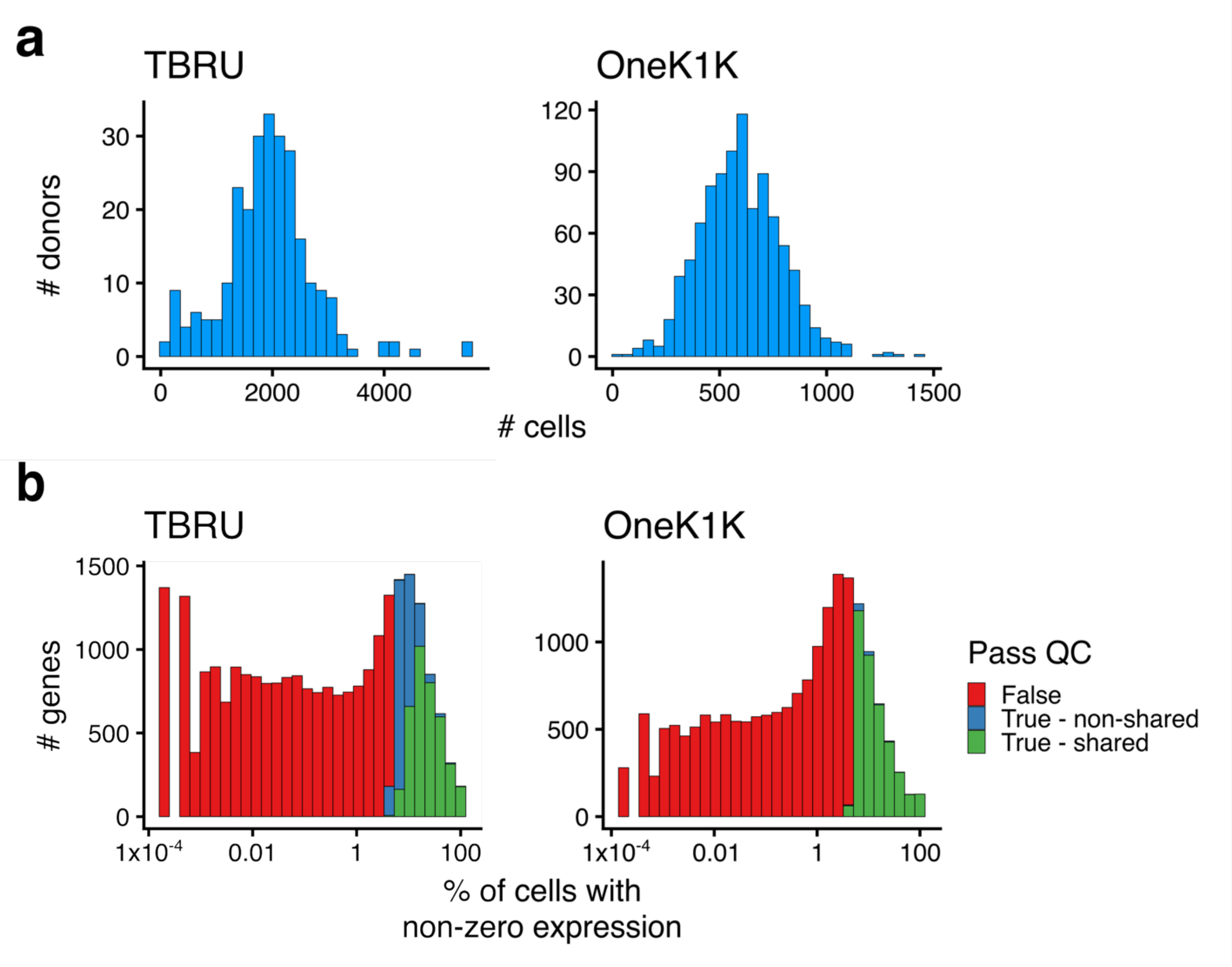


**Supplementary Figure 4 | Expression summary statistics for TBRU and OneK1K datasets. a)** Number of cells per donor in TBRU (left) and OneK1K (right) datasets. **b)** Distribution of autosomal genes by % of cells with non-zero expression. Genes passing QC (non-zero expression in >50% of donors and >5% of cells) were expressed in both datasets (green) or only one dataset (blue). Genes failed QC are shown in red

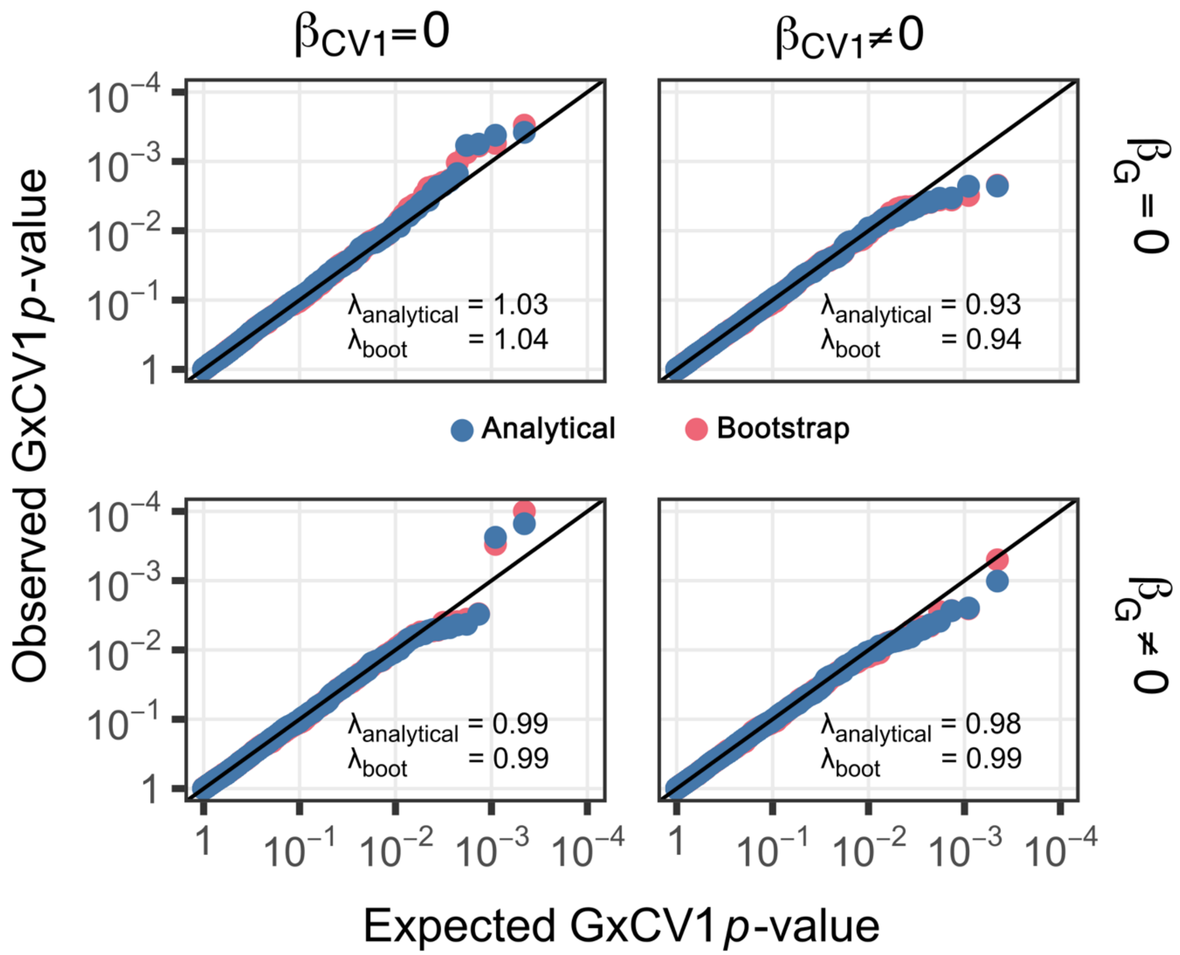


**Supplementary Figure 5 | *Dynema*'s calibration for testing interaction eQTL effects with CV1 under different null scenarios in the TBRU dataset.** *Dynema*'s analytical and bootstrap interaction-effect calibration under different scenarios generated by simulating the presence ($\beta = 0.2$) or absence ($\beta = 0$) of genotype (G) and/or cell state (CV1) main effects. In each scenario there is no simulated interaction between G and CV1. Each panel shows a Q-Q plot where each point corresponds to a lead variant-gene test (2,202 in total). The y-axis value is the *p*-value calculated *with Dynema* for the interaction between the lead variant ascertained through previous pseudobulk eQTL analysis and the CV1 cell state (i.e. $\beta_{GxCV1}$), and the x-axis is the *p*-value expected under a uniform null distribution. Statistical inflation is summarized with lambda values calculated for both analytical (blue) and bootstrap (red) p-values.

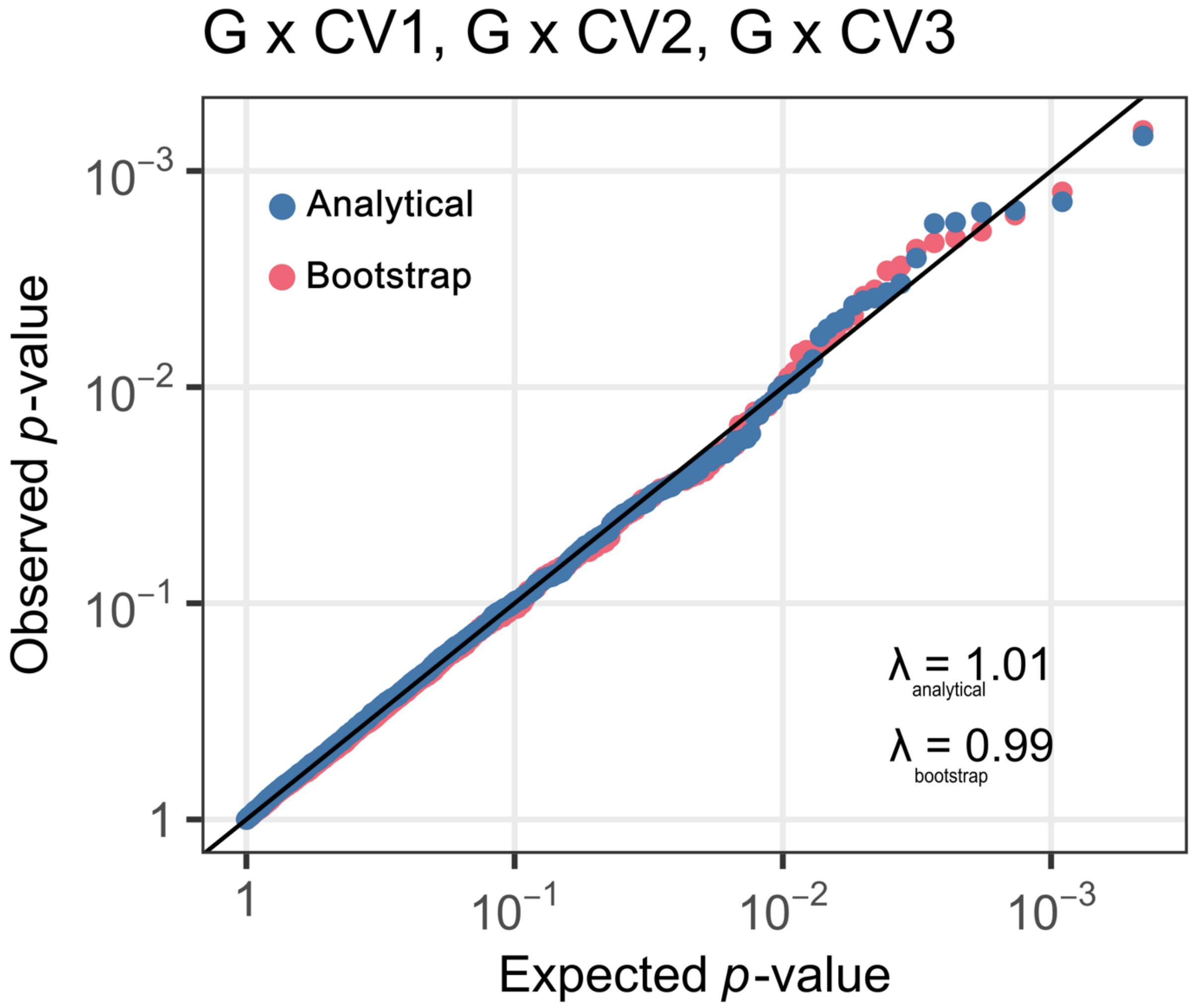


**Supplementary Figure 6 | *Dynema*'s calibration for testing multi-context interaction eQTL effects with CV1-3 under a null scenario in the TBRU dataset**. Q-Q plot showing the expected and observed *p*-values of *Dynema* via genotype permutation for 2,202 variant-gene pairs. Each point corresponds to a lead variant-gene test (2,202 in total). The y-axis value is the *p*-value calculated *with Dynema* for the multi-context interaction between the lead variant ascertained through previous pseudobulk eQTL analysis and CV1-3 cell states jointly, and the x-axis is the *p*-value expected under uniform null distribution. Statistical inflation is summarized with lambda values calculated for both analytical (blue) and bootstrap (red) *p*-values.

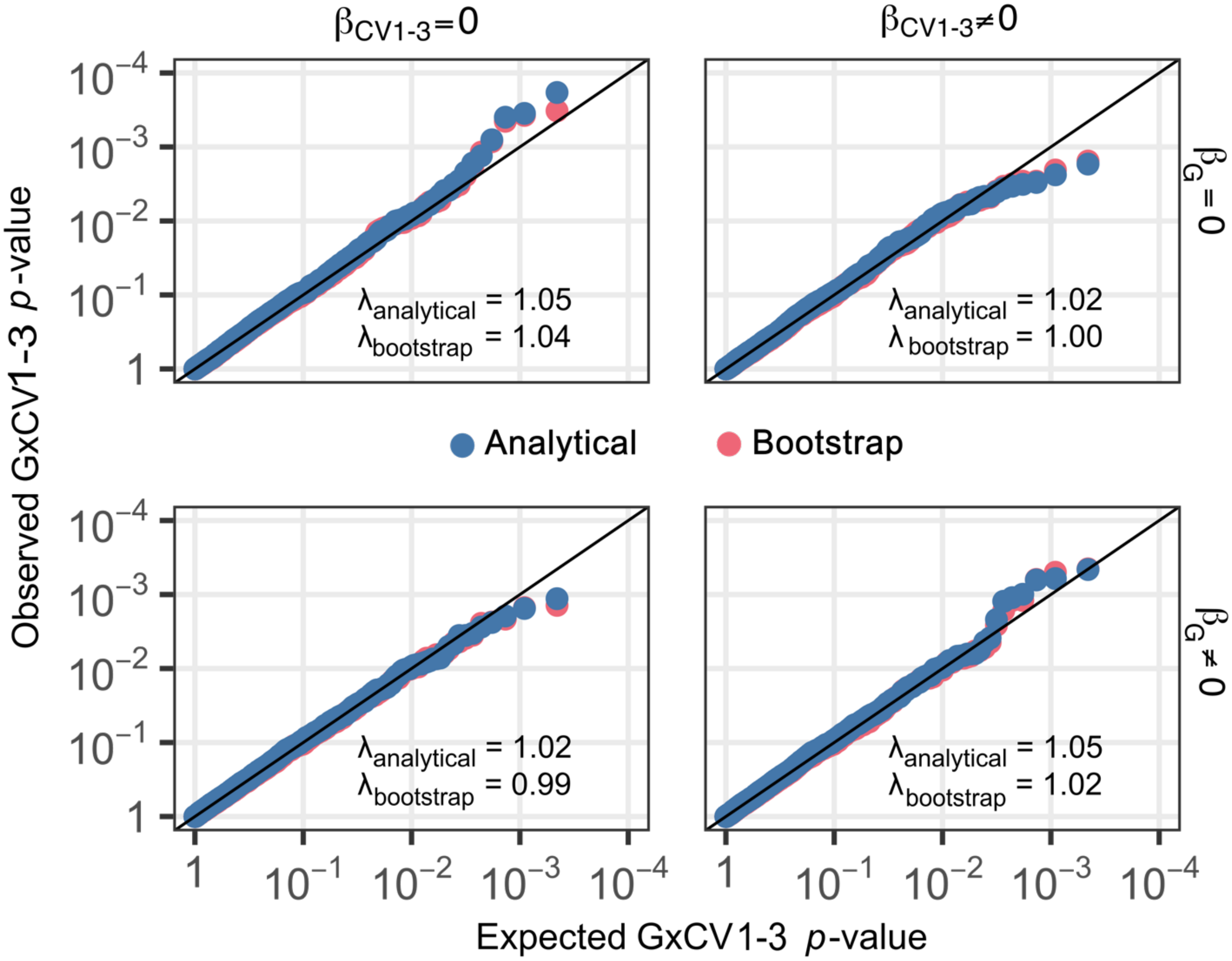


**Supplementary Figure 7 | *Dynema's* calibration for testing multi-context interaction eQTL effects with CV1-3 under different null scenarios in the TBRU dataset.** Calibration of Dynema's analytical and bootstrap interaction-effect under different scenarios generated by simulating the presence ($\beta$= 0.2) or absence ($\beta$= 0) of genotype (G) and/or presence (CV1-3 = 0.2) or absence (CV1-3 = 0) of cell state main effects. In each scenario there is no modeled interaction between G and CV1-3. Each panel shows a Q-Q plot where each point corresponds to a lead variant-gene test (2,202 in total). The y-axis value is the p-value calculated with *Dynema* for the multi-context interaction between the lead variant ascertained through previous pseudobulk eQTL analysis and CV1-3 cell states jointly, and the x-axis is the p-value expected under a uniform null distribution. Statistical inflation is summarized with lambda values calculated for both analytical (blue) and bootstrap (red) *p*-values.

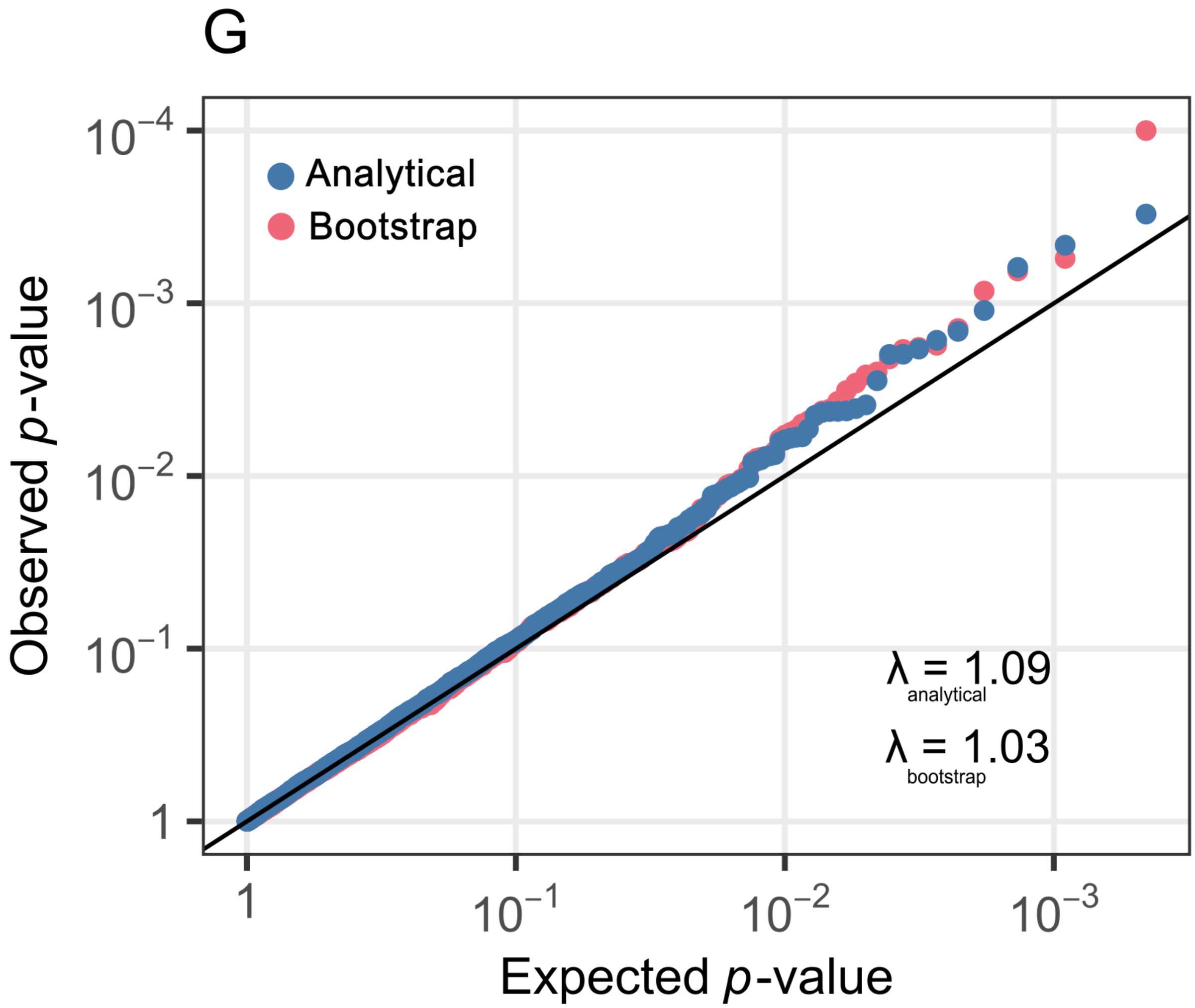


**Supplementary Figure 8 | *Dynema*'s calibration at testing main eQTL effects in the TBRU dataset**. Q-Q plot showing the expected and observed *p*-values of *Dynema* via genotype permutation for 2,202 variant-gene pairs. Each point corresponds to a lead variant-gene test (2,202 in total). The y axis value is the *p*-value calculated *with Dynema* for the genotype term, and the x axis is the p-value expected under a uniform null distribution. Statistical inflation is summarized with lambda values calculated for both analytical (blue) and bootstrap (red) *p*-values.

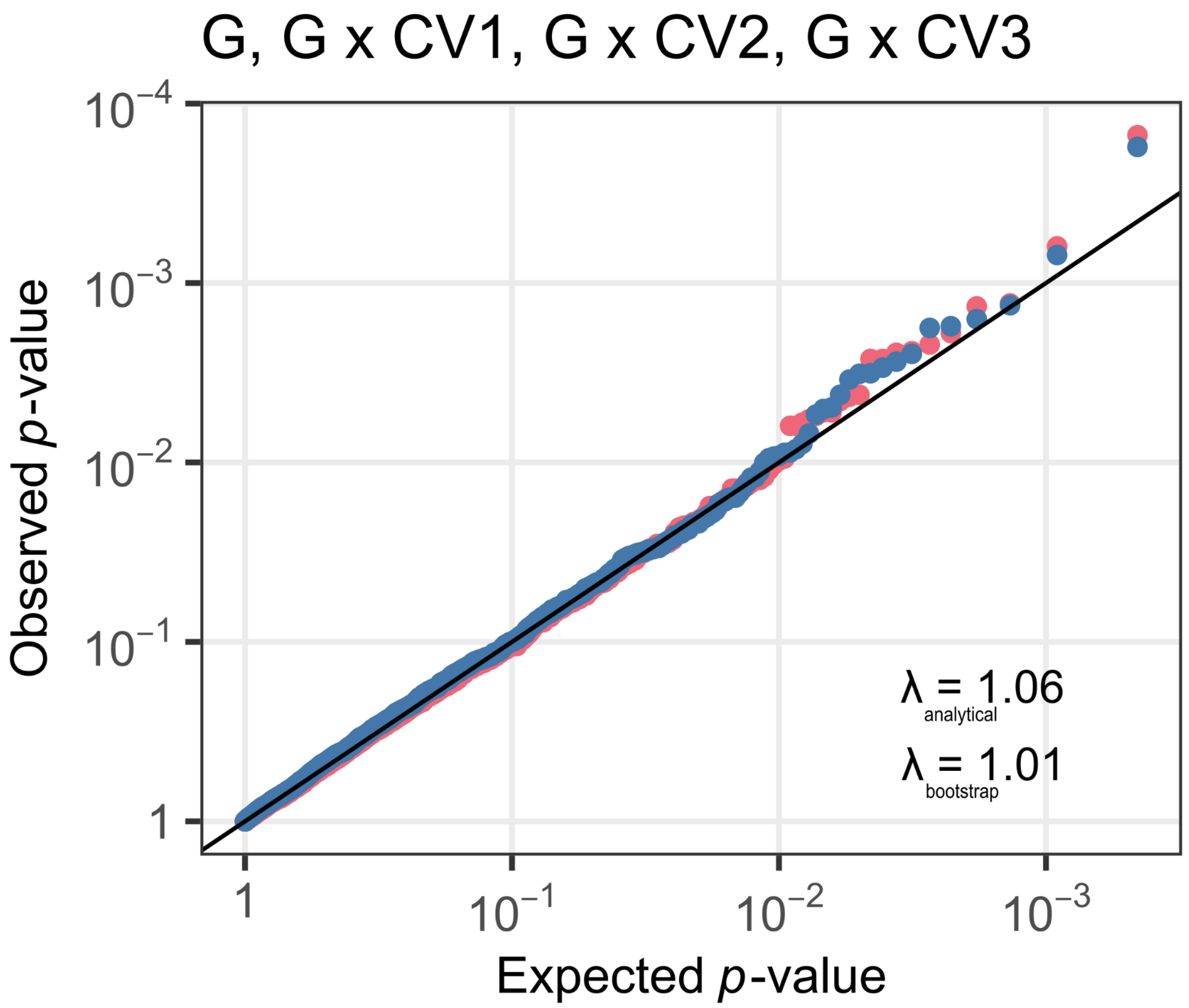


**Supplementary figure 9 | *Dynema*'s calibration for testing total eQTL effect with CV1-3 in the TBRU dataset**. Q-Q plot showing the expected and observed p-values of *Dynema* via genotype permutation for 2,202 variant-gene pairs. Under this null scenario, we plot expected and observed p-values for the total eQTL test capturing a genotype main effect and genotype interaction effects with each of the three CVs. Each point corresponds to a lead variant-gene test (2,202 in total). The y axis value is the p-value calculated *with Dynema* for the genotype main effect and multi-context interaction between the lead variant ascertained through previous pseudobulk eQTL analysis and CV1-3 cell states jointly, and the x axis is the *p*-value expected under a uniform null distribution. Statistical inflation is summarized with lambda values calculated for both analytical (blue) and bootstrap (red) *p*-values.

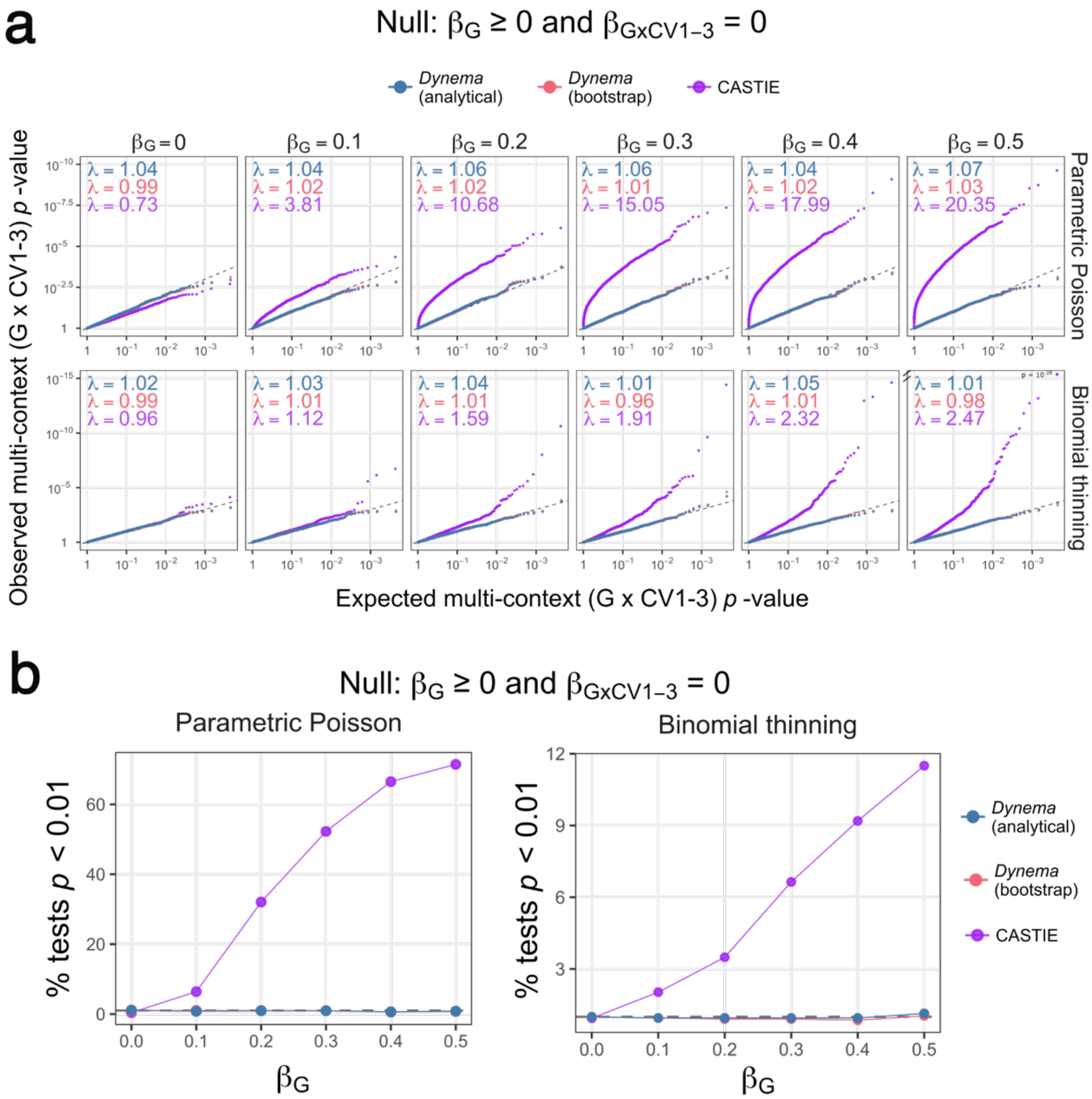


**Supplementary Figure 10 | Calibration of *Dynema* and CASTIE*'s* eQTL interaction statistics in the presence of main effects. a,** X and Y axes represent the expected and observed multi-context interaction (joint G x CV1-3) *p*-values respectively. Each column shows a different null scenario ($\beta_{GxCV1\text{-}3} = 0$) with increasing main effect from left to right. Each row corresponds to each of the simulation strategies: parametric Poisson (top) and binomial thinning (bottom). Each dot represents a variant-gene pair for *Dynema* (n=2,202) and CASTIE (n=2,122; 80 pairs failed computation with CASTIE, when gene names included non-alphanumeric characters). Expected *p*-values and lambda calculations were performed independently for each method to account for the difference in the number of tested variant-gene pairs. **b**, percent of significant tests at $p < 0.01$ in parametric Poisson (left) and binomial thinning (right) simulations. The X-axis shows the main-effect beta increasing from left to right. Y-axis shows the % of significant tests at $p < 0.01$. The expected percent of significant events at the null is 1% (dashed line).

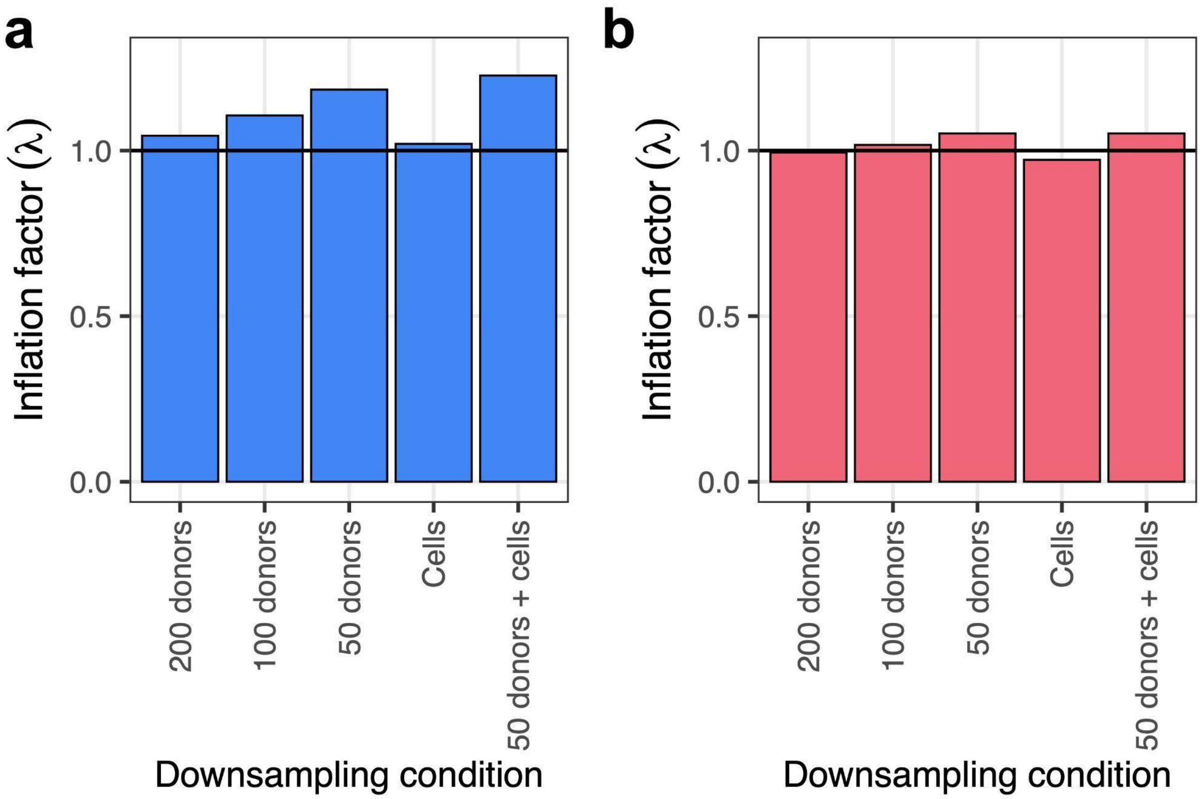


**Supplementary Figure 11 | *Dynema* calibration in downsampled conditions.** Inflation factor ($\lambda$) for **a)** *Dynema* analytical p-values and **b)** *Dynema* bootstrapped p-values, using downsampled gene expression data. We generated downsampled gene expression for each of the 2,202 SNP-gene pairs used for calibration analyses. For these analysis we either downsampled donors selecting a random subset of 50, 100, or 200 donors from the original 259 (first three bars), or (2) we considered all donors, and in half the donors we downsampled cells (selecting a random subset of 10%, 25%, or 50% of cells differently per donor, fourth bar), or (3) downsampled to 50 random donors, and in half of those donors we downsampled cells (selecting a random subset of 10%, 25%, or 50% of cells differently per donor, fifth bar).

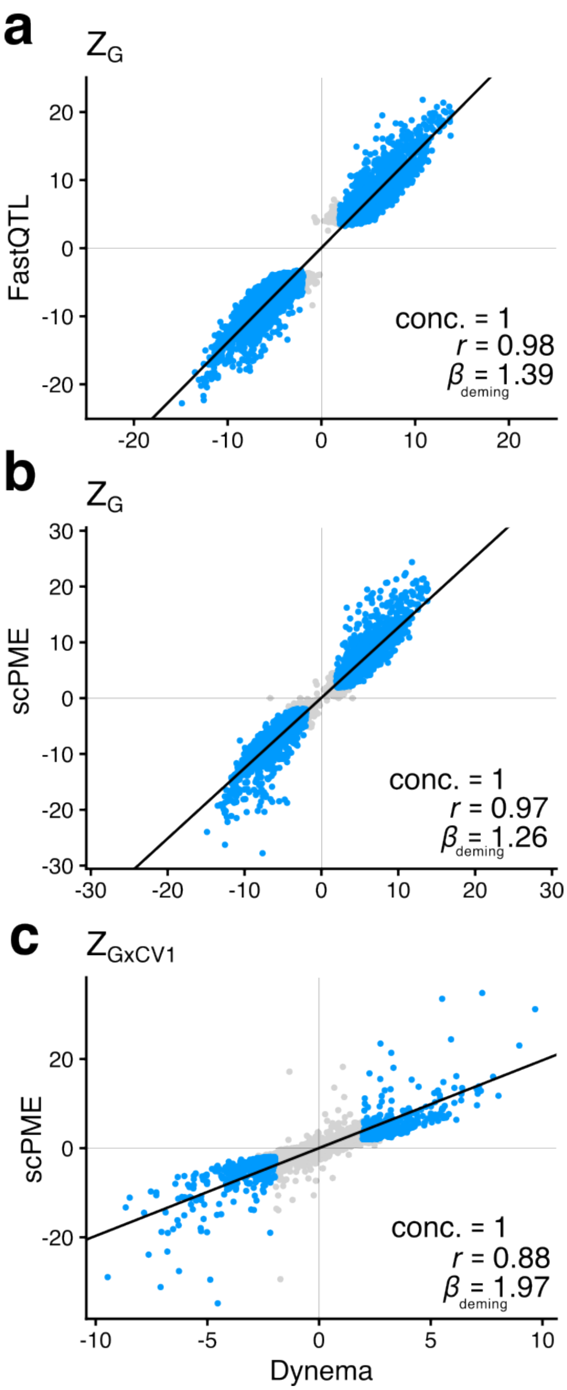


**Supplementary Figure 12 | *Dynema* recapitulates eQTL effects from FastQTL and scPME models**. Main effect eQTL z-score estimates from *Dynema* (x-axis) compared to **a,** FastQTL, or **b,** scPME (y-axis). **c.** GxCV1 interaction z-scores from *Dynema* (x-axis) compared to scPME (y-axis). Each point represents a variant-gene pair, and significant variant-gene pairs ($p$ < 0.05) across both plotted methods are shown in blue. Allelic directionality concordance, Pearson correlation, and slope from Deming regression are shown in the right-bottom are of each panel plot.

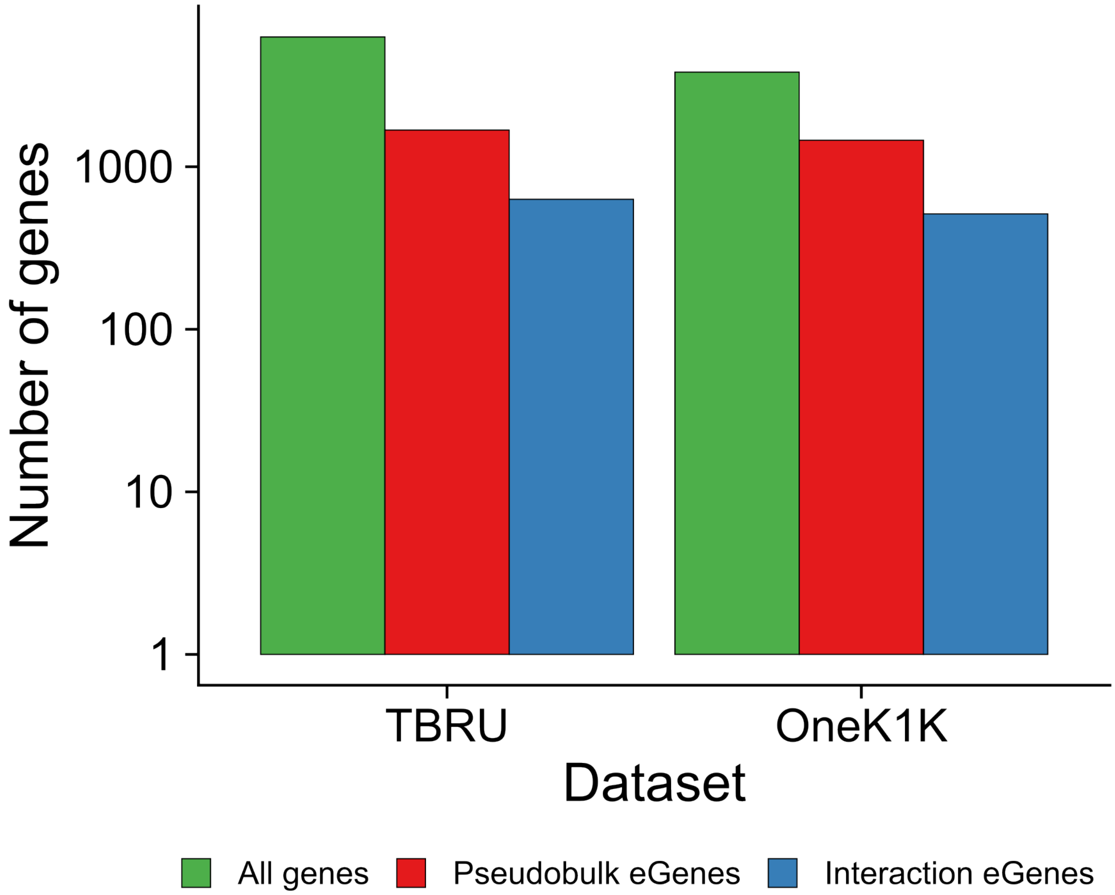


**Supplementary Figure 13 | *Dynema* identifies cell-state-dependent regulatory effects for pseudobulk eQTL variants.** The y-axis ($log_{10}$ scale) shows the number of genes tested (green bar), pseudobulk eGenes (blue bar), and pseudobulk eGenes with interaction eQTL effects identified with *Dynema* (red bar) in both TBRU and OneK1K (x-axis) by testing the lead main effect variant from pseudobulk for interaction with CV1-3.

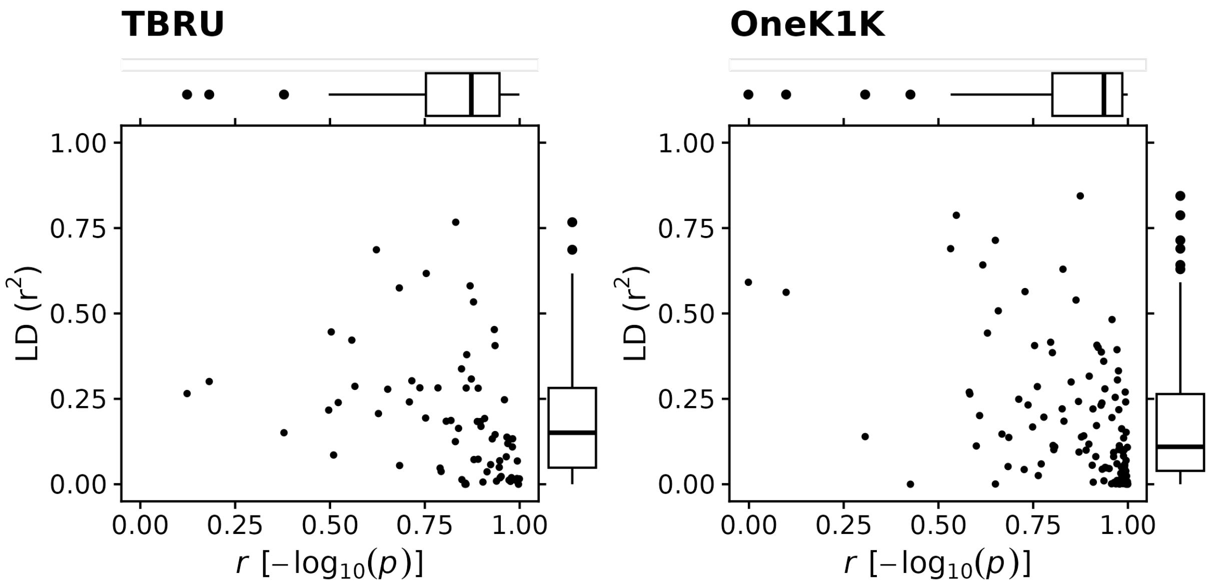


**Supplementary Figure 14 | Correlation of locus-wide *p*-values before and after conditioning vs LD between main and interaction lead effect variants.** Each dot represents a conditionally independent interaction eGene in TBRU (left) and OneK1K (right). x-axis shows the pairwise Pearson correlation of log-transformed interaction *p*-values before and after conditioning on the main effect lead variant across all variants within the same locus. y-axis shows the LD information ($r^2$) between the interaction and main effect lead variants for each locus. Most independent interaction effects from the main effect show high correlation and low LD as expected.

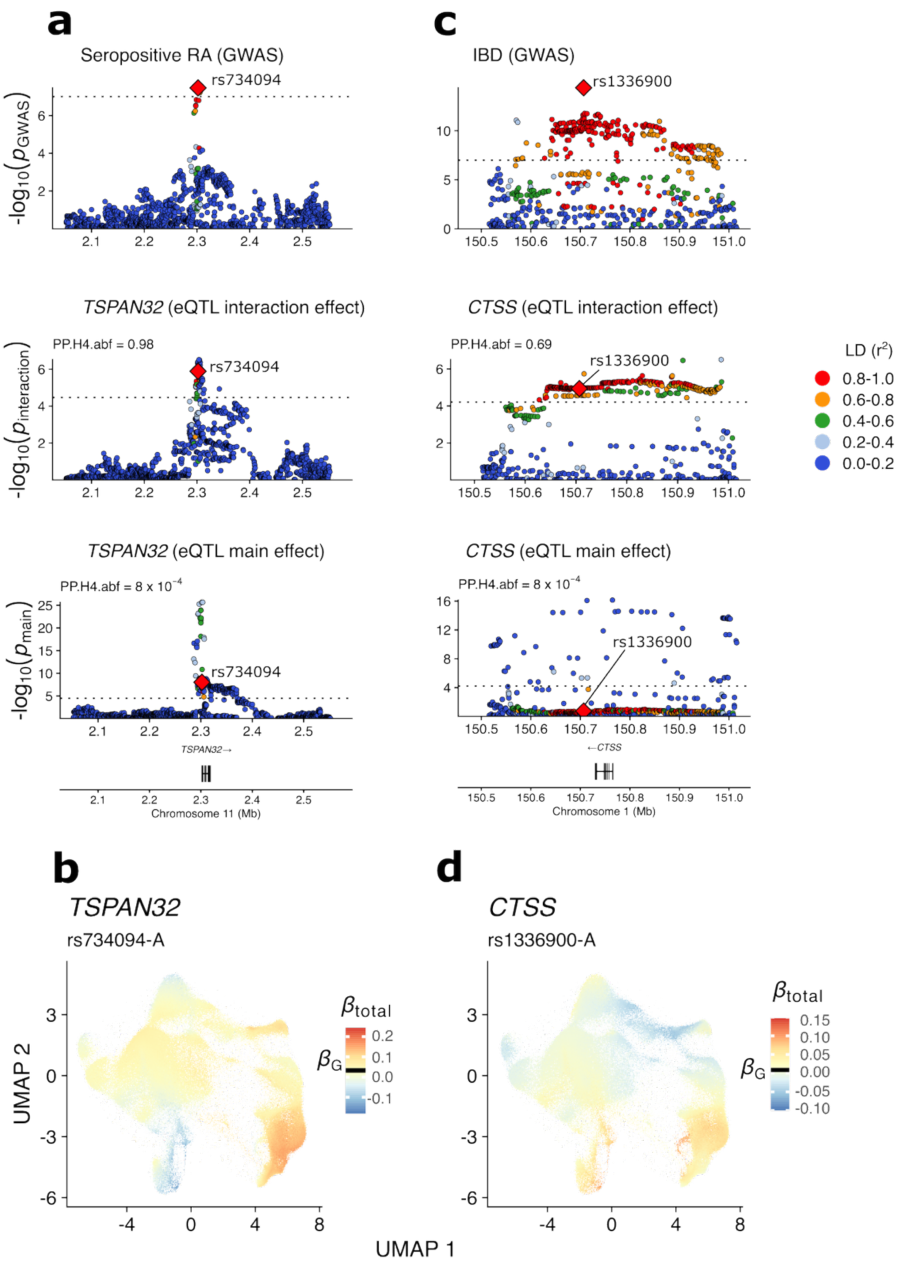


**Supplementary Figure 15 | Colocalization of autoimmune disease GWAS loci variant with eQTL multi-context interaction effect in OneK1K.** Locus zoom plots are shown for autoimmune disease GWAS (top), multi-context interaction effect (middle), and main effect (bottom) in OneK1K for **a,** *TSPAN32* and **c**, *CTSS* loci colocalizing with Rheumatoid Arthritis (RA) and Inflammatory Bowel Disease (IBD), respectively. The red diamond represents the lead GWAS disease variant in each of the two loci. LD information ($r^2$) of all variants with the lead GWAS is represented as color gradient. Posterior probability for common causal genetic signal between autoimmune disease and eQTL effects are shown for multi-context interaction (middle) and main (bottom) effects. **c-d**, UMAP plots with TSCE of lead interaction variants for *TSPAN32* (**b**) and *CTSS* (**d**). Color scales are marked with a dash to indicate the main effect size, $\beta_G$.
